# Bacterial-type plant ferroxidases tune local phosphate sensing in root development

**DOI:** 10.1101/2021.03.19.436157

**Authors:** Christin Naumann, Marcus Heisters, Wolfgang Brandt, Philipp Janitza, Carolin Alfs, Nancy Tang, Alicia Toto Nienguesso, Joerg Ziegler, Richard Imre, Karl Mechtler, Yasin Dagdas, Wolfgang Hoehenwarter, Gary Sawers, Marcel Quint, Steffen Abel

## Abstract

Fluctuating bioavailability of inorganic phosphate (Pi), often caused by complex Pi-metal interactions, guide root tip growth and root system architecture for maximizing the foraged soil volume. Two interacting genes in *Arabidopsis thaliana*, *PDR2* (P5-type ATPase) and *LPR1* (multicopper oxidase), are central to external Pi monitoring by root tips, which is modified by iron (Fe) co-occurrence. Upon Pi deficiency, the *PDR2-LPR1* module facilitates cell type-specific Fe accumulation and cell wall modifications in root meristems, inhibiting intercellular communication and thus root growth. LPR1 executes local Pi sensing, whereas PDR2 restricts LPR1 function. We show that native LPR1 displays specific ferroxidase activity and requires a conserved acidic triad motif for high-affinity Fe^2+^ binding and root growth inhibition under limiting Pi. Our data indicate that substrate availability tunes LPR1 function and implicate PDR2 in maintaining Fe homeostasis. LPR1 represents the prototype of an ancient ferroxidase family, which evolved very early upon bacterial colonization of land. During plant terrestrialization, horizontal gene transfer transmitted LPR1-type ferroxidase from soil bacteria to the common ancestor of Zygnematophyceae algae and embryophytes, a hypothesis supported by homology modeling, phylogenomics, and activity assays of bacterial LPR1-type multicopper oxidases.

## Introduction

Optimal plant growth exquisitely depends on edaphic resources. The pivotal role of inorganic phosphate (H_2_PO_4_^-^ or Pi) in metabolism, paired with its scarce bioavailability, render the mineral nutrient a strongly restrictive factor in terrestrial primary production (Lopez-Arredondo et al., 2014; Crombez et al., 2019). Insolubility of Pi salts and immobility of Pi complexed on clay or metal oxide minerals severely restrict P accessibility. Thus, plants actively seek and mine this crucial macroelement, but must concurrently navigate Pi-associated metal toxicities (Al, Fe) by adjusting root system architecture and modifying rhizosphere chemistry (Kochian et al., 2015; Abel, 2017; Gutierrez-Alanis et al., 2018). When challenged by Pi limitation, most dicotyledonous plants attenuate primary root extension and stimulate lateral root development for increasing the soil volume foraged by multiple root tips, which are the hotspots for Pi capture (Kanno et al., 2016). Root tips monitor heterogeneous Pi distribution (local Pi sensing) for guiding root development (Peret et al., 2014; Abel, 2017). In *Arabidopsis thaliana*, Pi deprivation rapidly attenuates root cell elongation (<2 h) in the transition zone and progressively inhibits cell division (<2 days) in the root apical meristem (RAM) (Müller et al., 2015; Balzergue et al., 2017). Persistent Pi starvation corrupts the stem-cell niche (SCN), which is followed by root growth arrest (Sanchez-Calderon et al., 2005; Ticconi et al., 2009). Notably, local Pi sensing depends on external Fe availability, which points to antagonistic biologic Fe-Pi interactions (Svistoonoff et al., 2007; Ticconi et al., 2009; Müller et al., 2015; Hoehenwarter et al., 2016; Balzergue et al., 2017; Dong et al., 2017; Godon et al., 2019; Wang et al., 2019).

Genetic approaches identified key components of root Pi-sensing (Abel, 2017; Gutierrez-Alanis et al., 2018; Crombez et al., 2019). A module of functionally interacting genes expressed in overlapping root cell types, *PHOSPHATE DEFICIENCY RESPONSE 2 (PDR2)*, *LOW PHOSPHATE RESPONSE 1 (LPR1)* and its close paralog *LPR2*, which plays a minor but additive role in the root response to Pi availability (Svistoonoff et al., 2007), encodes proteins of the secretory pathway and targets both cell elongation and cell division in Pi-deprived root tips. PDR2, the single orphan P5-type ATPase (AtP5A), functions in the endoplasmic reticulum (Ticconi et al., 2004; Jakobsen et al., 2005; Dunkley et al., 2006; Ticconi et al., 2009; Sorensen et al., 2015; Naumann et al., 2019) whereas LPR1, a multicopper oxidase (MCO) with presumed Fe^2+^-oxidizing activity, is localized to cell walls (Svistoonoff et al., 2007; Müller et al., 2015). On low Pi, the *PDR2-LPR1* module mediates LPR1-dependent, root cell type-specific Fe^3+^ accumulation in the apoplast, which correlates with ROS (reactive oxygen species) generation and callose deposition (Müller et al., 2015). While ROS formation promotes peroxidase-dependent cell wall stiffening in the transition zone (Balzergue et al., 2017), callose deposition interferes with cell-to-cell communication and thus inhibits RAM activity (Müller et al., 2015). LPR1-dependent root cell differentiation likely intersects with peptide and brassinosteroid signaling (Gutierrez-Alanis et al., 2017; Singh et al., 2018).

Current evidence points to LPR1 as a principal component of Fe-dependent Pi sensing. Upon Pi limitation, insensitive *lpr1* mutations cause unrestricted primary root extension by preventing Fe accumulation and callose deposition in root tips. Because loss of *LPR1* (or external Fe withdrawal) suppresses hypersensitive *pdr2* root phenotypes in low Pi, PDR2 restricts LPR1 function; however, the underlying mechanisms are unknown (Müller et al., 2015; Hoehenwarter et al., 2016; Naumann et al., 2019). Thus, the biochemical identity of LPR1 and the mechanism of LPR1 activation upon Pi deprivation need to be established. Here we determine the catalytic properties of purified native and mutant LPR1 variants and monitor *LPR1* expression in root tips. We show that *LPR1* encodes a prototypical, novel ferroxidase with high substrate (Fe^2+^) specificity and affinity. While *LPR1* is expressed in root meristems and is independent of *PDR2* function or Pi and Fe availability, LPR1-dependent root growth inhibition in limiting Pi is highly sensitive to low external Fe concentration. LPR1 substrate availability governs the local Pi deficiency response, whereas PDR2 maintains homeostasis of LPR1 iron reactants. Intriguingly, LPR1-like proteins, characterized by possessing in their active site an acidic triad motif and distinctive Fe^2+^-binding loop, are present in all extant land plants, in Zygnematophyceae algae, and in soil bacteria. Our phylogenetic and biochemical analyses support the hypothesis that LPR1-type ferroxidases evolved very early during bacterial land colonization and appeared in plants through horizontal gene transfer from Terrabacteria to the common ancestor of Zygnematophyceae and Embryophyta.

## Results

### LPR1 expression in root meristems is independent of PDR2 and external Pi supply

Analysis of *pLPR1^Col^::eGFP* expression in Pi-replete *A. thaliana* primary and secondary root tips revealed highest *pLPR1* promoter activity in the SCN and weaker GFP-derived fluorescence in proximal endodermal and cortical cells (Fig. 1a). We compared cell-specific *pLPR1* promoter activity in wild-type and *pdr2* roots upon transfer of Pi-replete seedlings (5 d-old) to +Pi or –Pi media for up to 7 d. The *pLPR1* expression domain and GFP intensity were similar between both genotypes on +Pi medium and did not change notably in the wild-type upon Pi deprivation. In *pdr2* root tips, *pLPR1::GFP* expression was maintained for at least 1 d on –Pi agar and thereafter ceased because of RAM disorganization (Fig. 1b). We observed a similar genotype-and Pi-independent *pLPR1::GFP* expression pattern in root tips of plants continuously grown on +Pi or –Pi medium for up to 4 d (Extended Data Fig. 1). Because gene expression and RAM activity rapidly respond to Pi deprivation (within 24 h) (Müller et al., 2015; Hoehenwarter et al., 2016, Balzergue et al., 2017), we analyzed steady-state mRNA levels in excised root tips 1 d after seedling transfer from Pi-replete to +Pi or –Pi media. Our data confirm that expression of *LPR1* and its functionally redundant sister gene, *LPR2*, is independent of external Pi supply or PDR2 function (Fig. 1c).

**Fig. 1.**
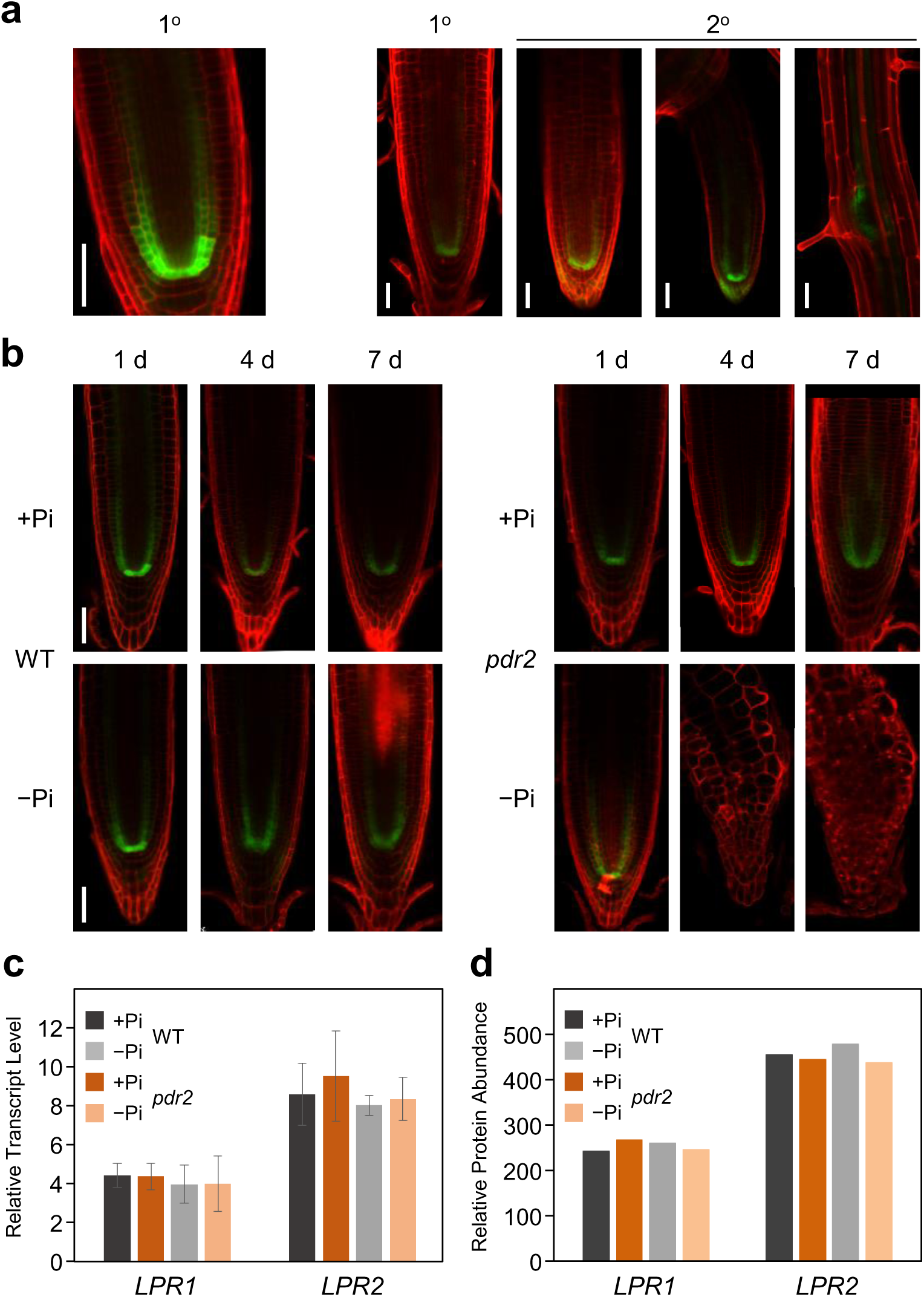
LPR1 expression in root meristems is independent of PDR2 and Pi availability. **a.** Expression of *pLPR1^Col^::GFP* in the SCN of transgenic wild-type primary (1°) and lateral (2°) root tips of seedlings continuously grown on +Pi medium for 6 d (left panel) or 12 d (right panels). Roots were counterstained with PI (red fluorescence), and GFP-derived fluorescence (green) was analyzed in primary and emerging lateral root tips. Shown are representative images (n≥7). Scale bars, 50 µm. **b.** Expression of *pLPR1^Col^::GFP* in primary root tips of transgenic wild-type (WT) and *pdr2* plants. Seeds were germinated for 5 d on +Pi agar prior to seedling transfer to +Pi or –Pi medium for up to another 7 d. Shown are representative images (n≥10). Scale bars, 50 µm. **c.** Relative transcript levels (normalized to *UBC9* expression) of *LPR1*, *LPR2*, and *PDR2* in excised WT and *pdr2* root tips (no significant differences, two-tailed Student’s *t*-test). Seeds were germinated on +Pi medium (5 d) and transferred to +Pi or –Pi medium. After 24 h, the gain in root tip growth was harvested for RNA preparations (± SD; n = 3). **d.** Relative protein abundance of LPR1 and LPR2 in excised root tips. Seeds were germinated on +Pi medium (5 d) and transferred to +Pi or –Pi medium. After 24 h, the gain in root tip growth was harvested for Tandem-Mass-Tag analysis. Shown are the sum of the normalized mass reporter intensities of LPR1 and LPR2 from one experiment.

To monitor LPR1 protein abundance, we generated peptide-specific anti-LPR1 antibodies that specifically recognize LPR1 in roots of overexpression (*pCaMV 35S::LPR1*) plants (Extended Data Fig. 2a, b). However, the antibodies failed to detect LPR1 in wild-type and *pdr2* root tips, even when profuse lateral root formation is stimulated to increase the number of root meristems, which express *pLPR1::GUS*. Likewise, attempts to enrich LPR1 by immuno-or chemical precipitation did not improve detection (Extended Data Fig. 2e-f). Eventually, using Tandem Mass Tag labeling (TMT) Mass Spectrometry, we detected with high confidence LPR1- and LPR2-derived peptides by quantitative proteomics on excised wild-type and *pdr2* root tips. Our data indicate that LPR1 and LPR2 abundance in root meristems does not depend on PDR2 function nor on external Pi status (Fig. 1d).

### Purified native LPR1 displays specific and high-affinity ferroxidase activity

The lack of evidence for Pi- or PDR2-dependent regulation of LPR1 expression prompted us to purify and characterize the LPR1 MCO enzyme. Because numerous attempts failed to express active, affinity-tagged recombinant LPR1, we purified native LPR1 to near homogeneity (monomeric protein of ∼70 kDa) from leaves of LPR1-overexpression plants (Fig. 2a). The three-step purification procedure involved differential ammonium sulfate precipitation followed by size exclusion and cation exchange chromatography, which yielded 2 μg LPR1 protein per gram leaf material (Extended Data Table 1). Immunoblot analysis, peptide sequencing by mass spectrometry, and ferroxidase activity assays verified the identity of LPR1, which was not detectable in identically prepared fractions from *lpr1lpr2* leaves (Extended Data Fig. 3, Extended Data Fig. 4).

**Fig. 2.**
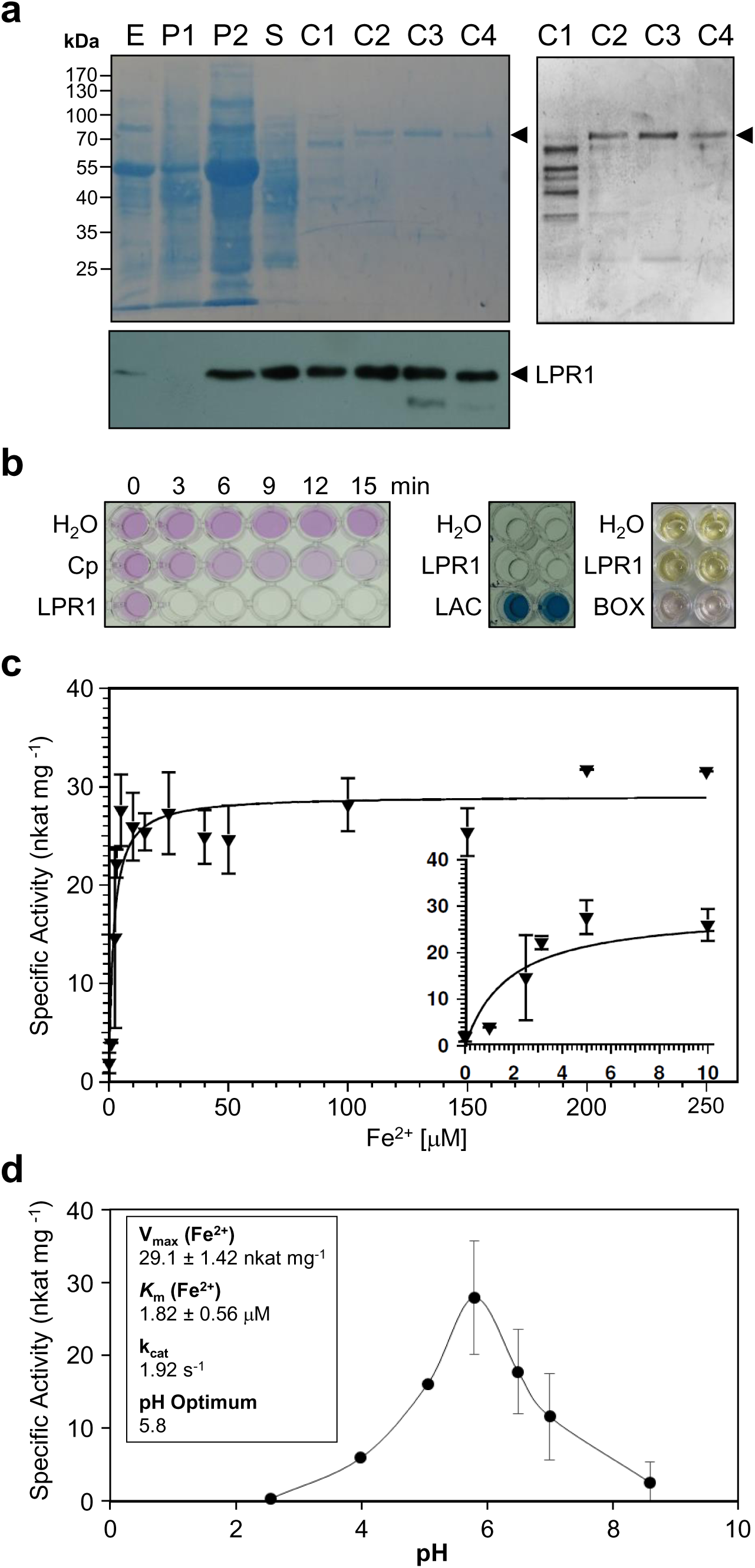
Purification and characterization of native LPR1 protein. **a.** Purification of native LPR1 from leaves of LPR1-overexpression plants. Proteins derived from all purification steps were separated by SDS-PAGE (upper left panel, Coomassie-stained gel, 20 µg protein per lane; upper right panel, silver-stained gel, 1 µg protein per lane) and transferred to membranes for immunoblot analysis using epitope-specific anti-LPR1 antibodies (lower panel). Protein fractions: (E) leaf extract; (P1) first ammonium sulfate (40% saturation) precipitation; (P2) second ammonium sulfate (40-80% saturation) precipitation; (S) size exclusion chromatography; (C1-C4) cation exchange chromatography. Arrow heads point to LPR1. **b.** Substrate specificity of LPR1. Left panel, discontinuous ferroxidase assay using 2 μg purified LPR1 and commercial ceruloplasmin (Cp). Pink color shows the substrate Fe^2+^-ferrozine complex (25 μM). Center panel, laccase assay (90 min) using purified LPR1 and commercial laccase from *Trametes versicolor* (LAC). The colorless substrate (500 μM ABTS) is oxidized to a colored product. Right panel, bilirubin oxidase assay (15 min, 30 μM bilirubin) comparing purified LPR1 and commercial bilirubin oxidase from *Myrothecium verrucaria* (BOX). **c.** Fe^2+^ concentration-dependent (0-250 μM) ferroxidase activity of purified, native LPR1 protein (n=3). Inset: LPR1 activity for low Fe^2+^ concentration range (0-10 μM). **d.** pH optimum of LPR1 ferroxidase activity in 0.1 M Na-acetate buffer (±SD, n=3). Inset: kinetic constants of LPR1.

The diverse MCO protein family comprises phenol oxidases (laccases), bilirubin oxidases, ascorbate oxidases, and metal oxidases such as ferroxidases from yeast (Fet3p) or humans (ceruloplasmin) (Graff et al., 2020). We previously reported elevated ferroxidase activity in root extracts of LPR1-overexpression plants (Müller et al., 2015). Using purified native LPR1 and the ferrozine assay, we determined kinetic parameters of its ferroxidase activity. LPR1 exhibited a typical Michaelis-Menten saturation kinetics for Fe^2+^ oxidation, which revealed an apparent *K*_m_ value of 1.8 μM, a V_max_ value of 29 nkat mg^-1^, and a turnover frequency *k*_cat_ of 1.9 s^-1^ (Fig. 2b-d). LPR1 displayed highest ferroxidase activity at pH 5.8, a value consistent with LPR1 function in the apoplast (Müller et al., 2015). In addition to its ferroxidase activity, we tested LPR1 for laccase activity with ABTS (2,2’-azino-bis[3-ethylbenzothiazoline-6-sulfonic acid]) as the substrate, and for ascorbate, bilirubin and manganese oxidase activities (Fig. 2b, Extended Data Fig. 5). The inability to oxidize the four MCO substrates indicates specificity of LPR1 for ferrous iron. Because proteins of the secretory pathway are often N-glycosylated, we treated purified LPR1 with a mixture of deglycosylating enzymes. We did not obtain evidence for LPR1 glycosylation or phosphorylation (Extended Data Fig. 4b-d). Peptide sequencing by mass spectrometry of purified native LPR1 indicated absence of both types of posttranslational modifications; however, ectopic overexpression of LPR1 in leaves may have overwhelmed and masked detectable LPR1 protein modifications.

### Local Pi sensing requires the Fe-binding site on LPR1

We previously derived a structural model of LPR1 based on its presumed function in Fe homeostasis and distant similarity to Fet3p, which suggested presence of an Fe^2+^-binding (Müller et al., 2015). We independently employed the YASARA package to identify by PSI-BLAST iterations and PDB searches high-scoring templates for LPR1 homology modeling. The five top-most ranking templates are all crystal structures of the spore-coat protein A (CotA), a bacterial (*Bacillus subtilis*) MCO laccase (Enguita et al., 2004). The refined LPR1 model reveals the hallmarks of MCO enzymes, i.e. the mononuclear T1 Cu site and the multinuclear T2/T3 Cu cluster, and it further supports the presence of an acidic triad (E269, D370, and D462) predicted to compose the Fe^2+^ binding site (Fig. 3a-c).

**Fig. 3.**
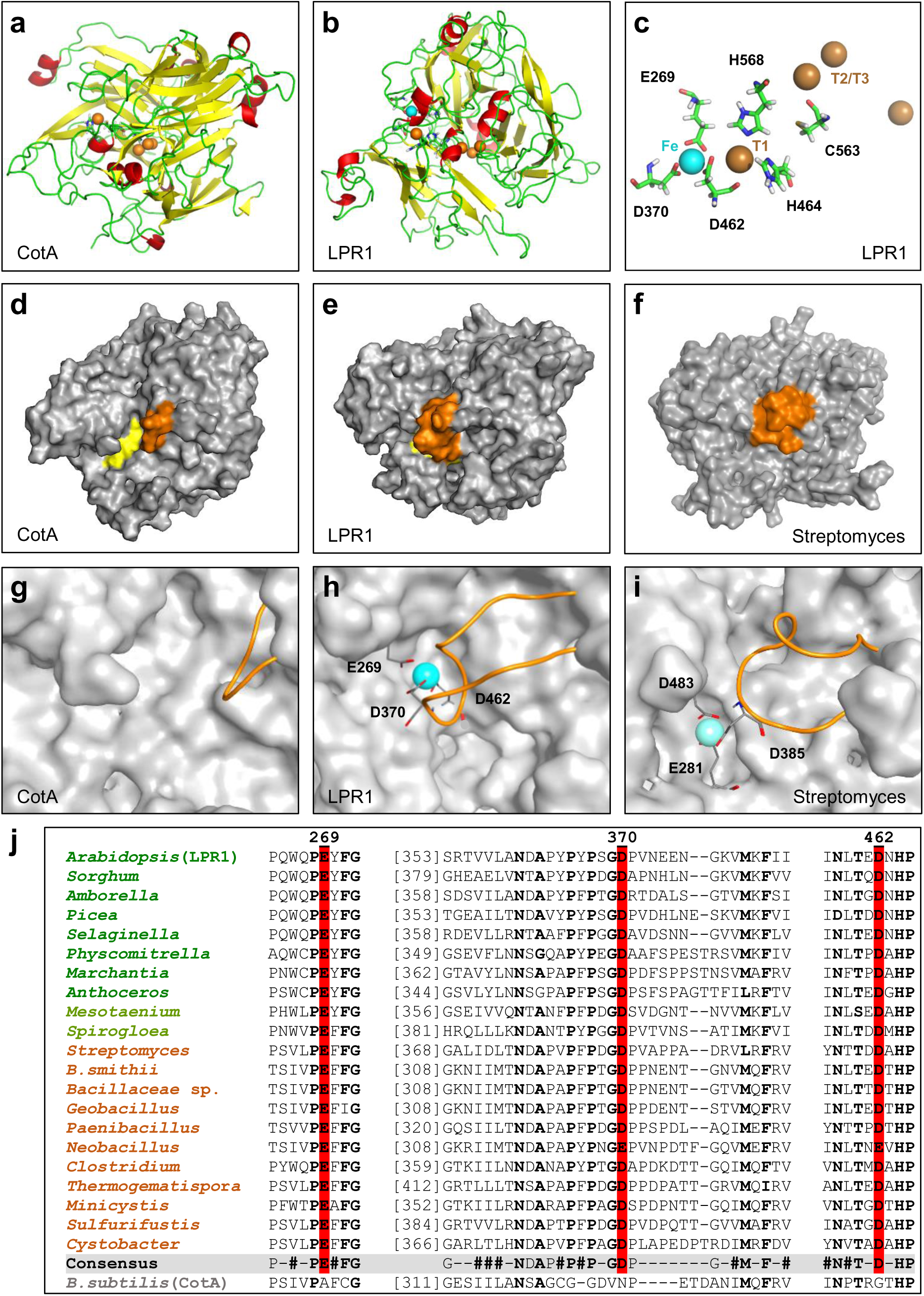
Homology models of LPR1 and related proteins. **a-c.** The 3D structure of CotA (**a**), an MCO laccase from *Bacillus subtilis* (PDB ID: 4AKP) was used as a template to model the tertiary structure of LPR1 (**b**), which accommodates the predicted four catalytic Cu sites (orange spheres) and the bound Fe^2+^ cation (blue sphere). The predicted LPR1 Fe^2+^ substrate-binding site (E269, D370 and D462), the proximal mononuclear T1 Cu site (H464, C563 and H568) and distal trinuclear T2/T3 Cu cluster are shown in (**c**). **d-i.** Surface representations of the experimental CotA structure (**d, g**), the LPR1 homology model (**e, h**), and the homology model of the LPR1-like MCO protein from *Streptomyces clavuligerus* (**f, i**). Overview structures (**d-f**) highlight the substrate-binding pocket (yellow) and adjacent loop (orange) on each protein model. The magnified substrate binding sites (**g-i**) depict the adjacent loop (orange cartoon), the acidic triad (sticks), and the Fe^2+^ substrate (blue sphere). **j.** Alignment of conserved sequence motifs flanking each residue of the acidic triad (highlighted in red) relative to the Fe-binding site on *Arabidopsis* LPR1 (E269, D370, D462). The central acidic residue (D370) is located in a variable linker sequence terminated by conserved border residues (see consensus). Aligned are the three acidic triad motifs of select LPR1-like MCOs of plants (green), Zygnematophycea (light green) and Terrabacteria (brown). Corresponding motifs in CotA are shown below.

We generated by site-directed mutagenesis cDNAs for expressing *in planta* LPR1 variants with single or multiple amino acid substitutions in the presumed Fe^2+-^binding pocket (E269A, D370A, D462A, and combinations thereof) or proximal T1 Cu site (H464A, H568A, and C563A). LPR1 expression (immunoblot analysis) and specific ferroxidase activity were determined in extracts from transiently transfected tobacco (*Nicotiana benthamiana*) leaf discs. Expression of wild-type LPR1 (*p35S::LPR1^WT^*) resulted in the highest specific ferroxidase activity of all plasmids tested (Fig. 4a). The LPR1^WT^ leaf extract exhibited typical Michaelis-Menten saturation kinetics for Fe^2+^ oxidation and revealed an apparent *K*_m_ value of 3.6 μM (Fig. 4a, inset), which is similar to the value obtained with purified wild-type LPR1 (Fig. 2c).

**Fig. 4.**
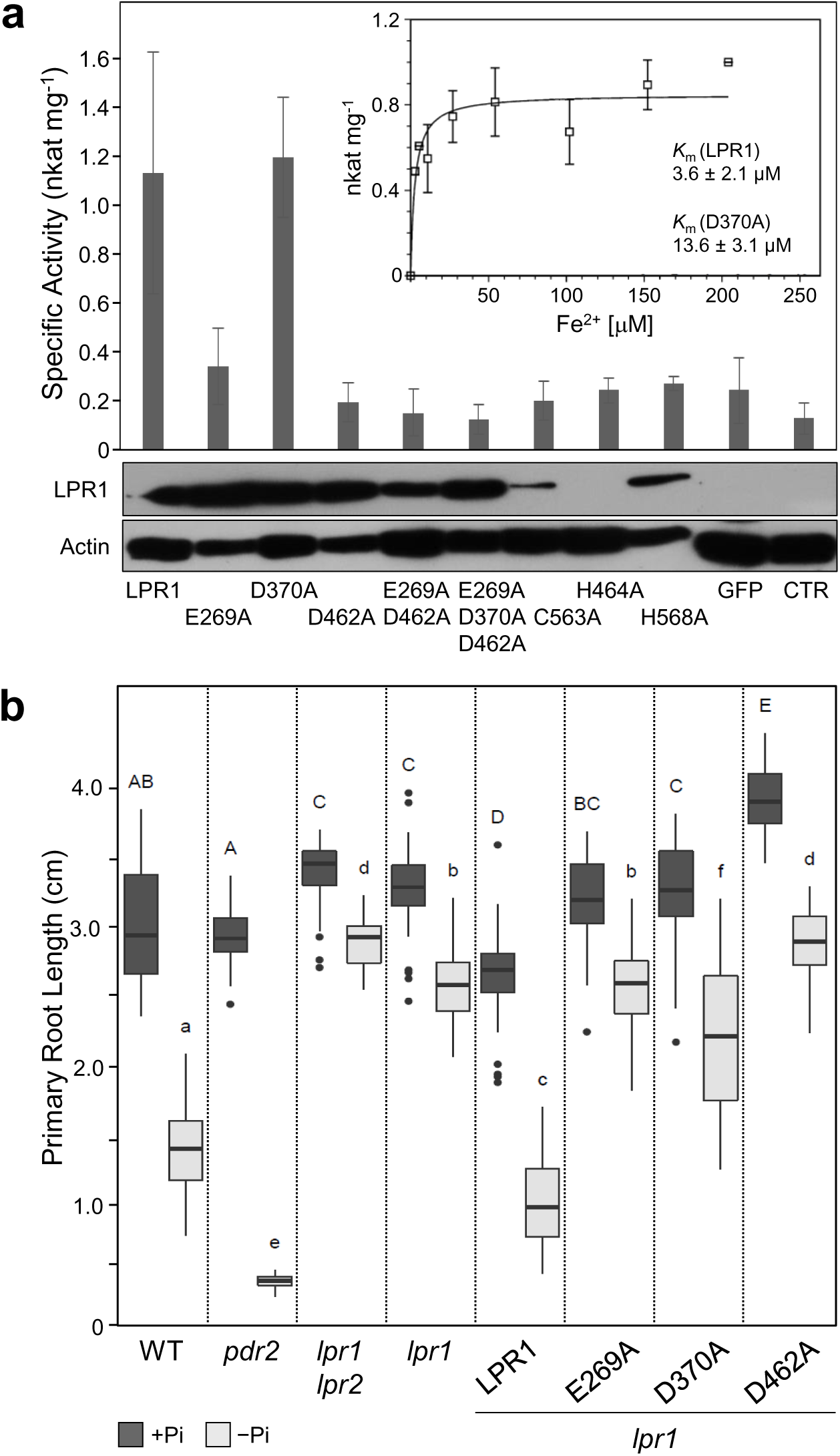
The Fe^2+^ binding site is required for LPR1 ferroxidase activity and local Pi sensing. **a.** Variants of *LPR1* cDNAs were generated by site-directed mutagenesis and the encoded proteins transiently expressed in tobacco leaves (*N. benthamiana*) under the control of the *CaMV 35S* promoter. Leaf discs were harvested 4 d after infiltration and protein extracts were prepared. Specific ferroxidase activity was determined (±SE; n≥3) for leaf discs overexpressing wild-type LPR1 protein or mutant LPR1 variants with single or multiple amino acid substitutions in the predicted Fe^2+^ binding site (E269A, D370A, D462A, and combinations thereof), or in the predicted mononuclear T1 Cu site (H464A, H568A, and C563A). Controls included infiltrations with buffer (CTR) or *p35S::GFP* plasmid DNA (GFP). Inset: Specific ferroxidase activity of leaf disc extracts overexpressing *p35S::LPR1* as a function of increasing Fe^2+^ substrate concentration (n=3). Lower panel: Protein extracts of leaf discs (12.5 µg) overexpressing the indicated LPR1 variants were used for immunoblot analysis with epitope-specific anti-LPR1 antibodies and monoclonal anti-actin (plant) antibodies. **b.** Seeds of the indicated genotypes were germinated for 5 d on +Pi agar containing 25 μM Fe prior to transfer to +P or –Pi medium (25 μM Fe). Extension of the primary root axis was recorded 4 d after transfer (±SD, n=27-36 seedlings). The following genotypes were tested: wild-type (WT), *pdr2*, *lpr1lpr2*, *lpr1*, as well as overexpression of *p35S::LPR1*, *p35S::LPR1^E269A^*, *p35S::LPR1^D370A^*, and *p35S::LPR1^D462A^* in *lpr1* plants. Box plots show medians and interquartile ranges of total root lengths; outliers (greater than 1.5× interquartile range) are shown as black dots. Different letters (+Pi: capital, -Pi: lower case) denote statistical differences in the respective condition at *P* < 0.05 as assessed by one-way ANOVA and Tukey’s honestly significant difference (HSD) posthoc test.

Amino acid substitutions in the acidic triad of the predicted Fe^2+^ binding site impaired specific ferroxidase activity to varying extents. While LPR1^D370A^ expression and specific activity were comparable to LPR1^WT^, leaf extracts expressing the LPR1^D370A^ variant showed an almost 4-fold higher *K*_m_-value for Fe^2+^ (13.6 μM) when compared to LPR1^WT^ leaf extracts (Fig. 4a). On the other hand, expression of LPR1^E269A^ showed an ∼80% reduction of the LPR1^WT^ ferroxidase activity measured above background, and expression of LPR1^D462A^ revealed an almost complete loss of ferroxidase activity, which was similar to that of control transfections. Expression of LPR1 variants with multiple amino acid substitutions, LPR1^E269A, D462A^ and LPR1^E269A, D370A, D462A^ did not increase specific ferroxidase activity above the background level, which was also observed for leaf extracts transfected with plasmids encoding LPR1 variants with a disrupted mononuclear T1 Cu site, i.e. LPR1^H464A^, LPR1^H568A^, and LPR1^C563A^ (Fig. 4a). However, the T1 Cu site mutant variants were noticeably less abundant, or undetectable (LPR1^H464A^), based on immunoblot analysis. Low protein abundance suggests LPR1 protein instability because Cu is a co-factor for both MCO activity and proper MCO protein folding.

To verify some of the results by *in planta* complementation, we generated *lpr1* plants expressing LPR1^WT^, LPR1^E269A^, LPR1^D370A^ and LPR1^D462A^ variants under control of the *pCaMV 35S* promotor. We compared primary root extension of the transgenic lines to wild-type and *pdr2*, as well as to *lpr1lpr2* and *lpr1* seedlings upon transfer from Pi-replete conditions (5 d) to +Pi or –Pi agar medium for 4 d (Fig. 4b). While overexpression of LPR1 restored primary root growth inhibition of insensitive *lpr1* seedlings on –Pi agar, overexpression of variants LPR1^E269A^ and LPR1^D462A^ did not significantly reduce the long root phenotype of *lpr1* seedlings on low Pi medium. We noticed a weak, albeit poorly, complementing effect for the LPR1^D370A^ variant (Fig. 4b). Thus, these data are largely consistent with the ferroxidase activity assays using tobacco leaves (Fig. 4a).

Using the LPR1 purification protocol, we prepared LPR1^E269A^, LPR1^D370A^ and LPR1^D462A^ variants for studying their kinetic parameters (Extended Data Fig. 6). We obtained low amounts of nearly pure LPR1^E269A^ and LPR1^D462A^ variants (detectable by immunoblot analysis), but we failed to prepare LPR1^D370A^. However, the final fractions of all LPR1 variant preparations did not display any ferroxidase activity, which points to a compromised Fe^2+^-binding pocket. In summary, our structure-function analysis of LPR1 firmly demonstrates a requirement of the predicted Fe^2+^-binding site for LPR1 ferroxidase activity (and possibly protein stability), as well as for LPR1-dependent Pi sensing by root meristems.

### Fe availability tunes LPR1-dependent Pi sensing

Because *LPR1* expression and LPR1 abundance do not noticeably respond to external Pi supply (Fig. 1), we tested in nutrient shift experiments whether LPR1 substrate availability governs local Pi sensing. We transferred Pi-replete seedlings (5-d-old) to +Pi or –Pi agar medium supplemented with increasing Fe concentrations (0-1000 μM) and monitored primary root extension for up to 4 d. Upon transfer to Fe-supplemented +Pi media, wild-type, *lpr1lpr2*, and *pdr2* seedlings displayed a similar daily extension of the primary root, which was not greatly altered by the addition of up to 200 μM Fe. Higher Fe concentrations (500-1000 μM) strongly inhibited primary root growth irrespective of the genotype, which is likely caused by general Fe toxicity (Reyt et al., 2015; Zhang et al., 2018) (Extended Data Fig. 7).

Seedling transfer from +Pi+Fe control agar to –Pi medium without iron (–Pi –Fe) does not significantly inhibit primary root extension of all genotypes tested, which we previously reported (Müller et al., 2015). However, we observed striking genotype-dependent differences in root growth inhibition upon transfer to –Pi media supplemented with increasing Fe (Fig. 5). Intriguingly, wild-type roots displayed a triphasic growth response to increasing Fe supply (Fig. 5a). Low Fe concentrations (2.5-25 μM, phase I) strongly reduced primary root extension (by 60%), whereas intermediate Fe availability (50-100 μM, phase II) was less inhibitory (ca. 30%). Higher Fe supply (>100 μM, phase III) caused a gradual and pronounced inhibition of primary root growth, which was also observed on +Pi medium supplemented with high (>500 μM) Fe concentrations (Extended Data Fig. 7a). Interestingly, *lpr1lpr2* seedlings did not display the triphasic Fe dose response in Pi deficiency (Fig. 5b). Primary root extension was insensitive to low Fe for up to 50 μM; however, *lpr1lpr2* root growth was gradually inhibited by higher Fe supply (100-1000 μM) in a similar fashion as wild-type roots (Fig. 5a, b). On the other hand, Pi-deprived *pdr2* seedlings displayed, as the wild-type, strong inhibition of primary root growth on gradually increasing, low Fe concentrations (2.5-25 μM). While root growth inhibition was maximal at 25 μM Fe (by 85%), higher concentrations (50-1000 μM Fe) neither rescued nor intensified *pdr2* root growth inhibition on –Pi media (Fig. 5c, d).

**Fig. 5.**
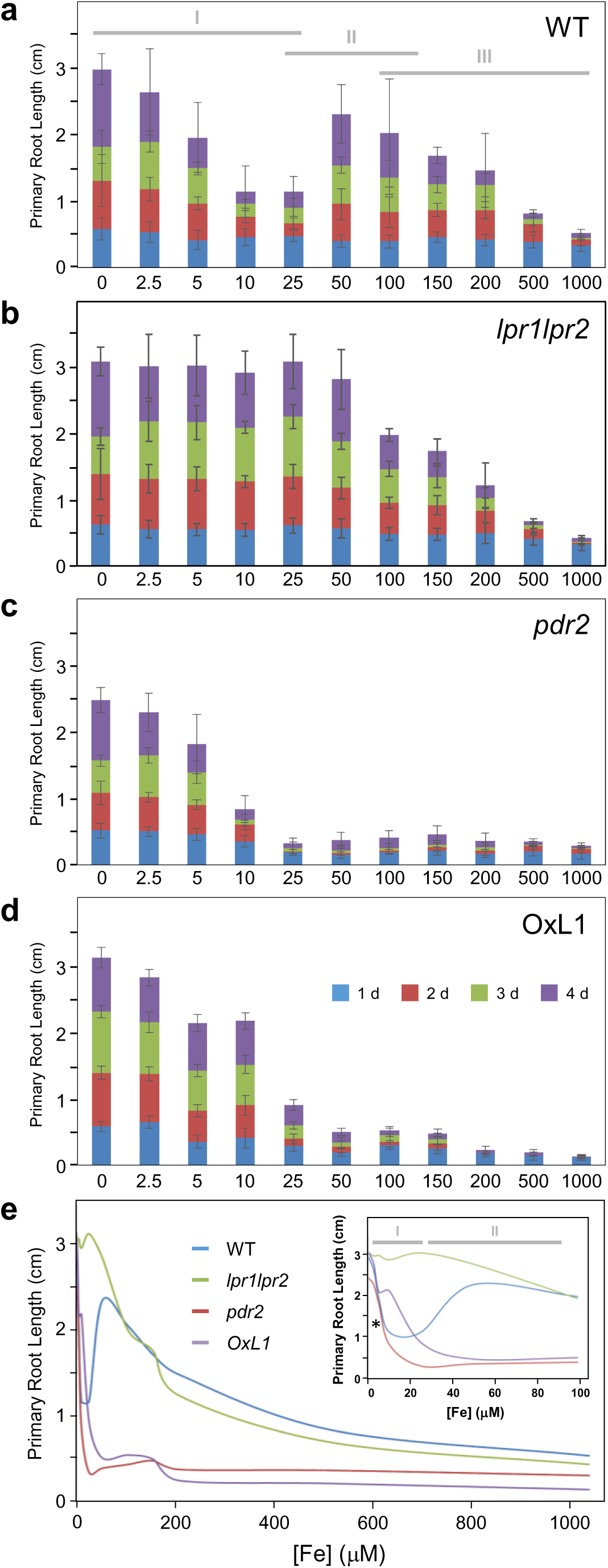
Iron-dependent inhibition of primary root growth upon Pi deprivation. **a-d.** Seeds of wild-type (WT), *lpr1lpr2*, *pdr2*, and LPR1-overexpression (OxL1) lines were germinated for 5 d on +Pi agar prior to transfer to –Pi media supplemented with increasing iron (Fe^3+^-EDTA) concentrations. Gain of primary root extension was measured daily for up to 4 d after transfer and plotted for each genotype (±SD; n≥50). The three ranges (I-III) of the Fe dose response curve of the wild-type are indicated in (**a**). **e.** Trend lines of Fe-dependent primary root growth on –Pi media from 0-1000 μM Fe. Inset: Trend lines for the low Fe concentrations (0-100 μM). Asterisk: *K*_m_ (Fe^2+^) of LPR1.

It is important to point out that the apparent *K*_m_ value of LPR1 (2-3 μM Fe^2+^) corresponds well with the first inhibition phase of primary root extension on low Fe (0-10 μM) under Pi limitation (Fig. 5e). This suggests that LPR1 ferroxidase activity and function in local Pi sensing are primarily determined by substrate availability. This proposition is supported by unaltered *pLPR1* and *pPDR2* promoter activities and by stable steady-state *LPR1* and *PDR2* mRNAs levels in wild-type root tips upon seedling transfer to –Pi agar supplemented with increasing (0-1000 μM) Fe concentration (Fig. 6a, Extended Data Fig. 8).

**Fig. 6.**
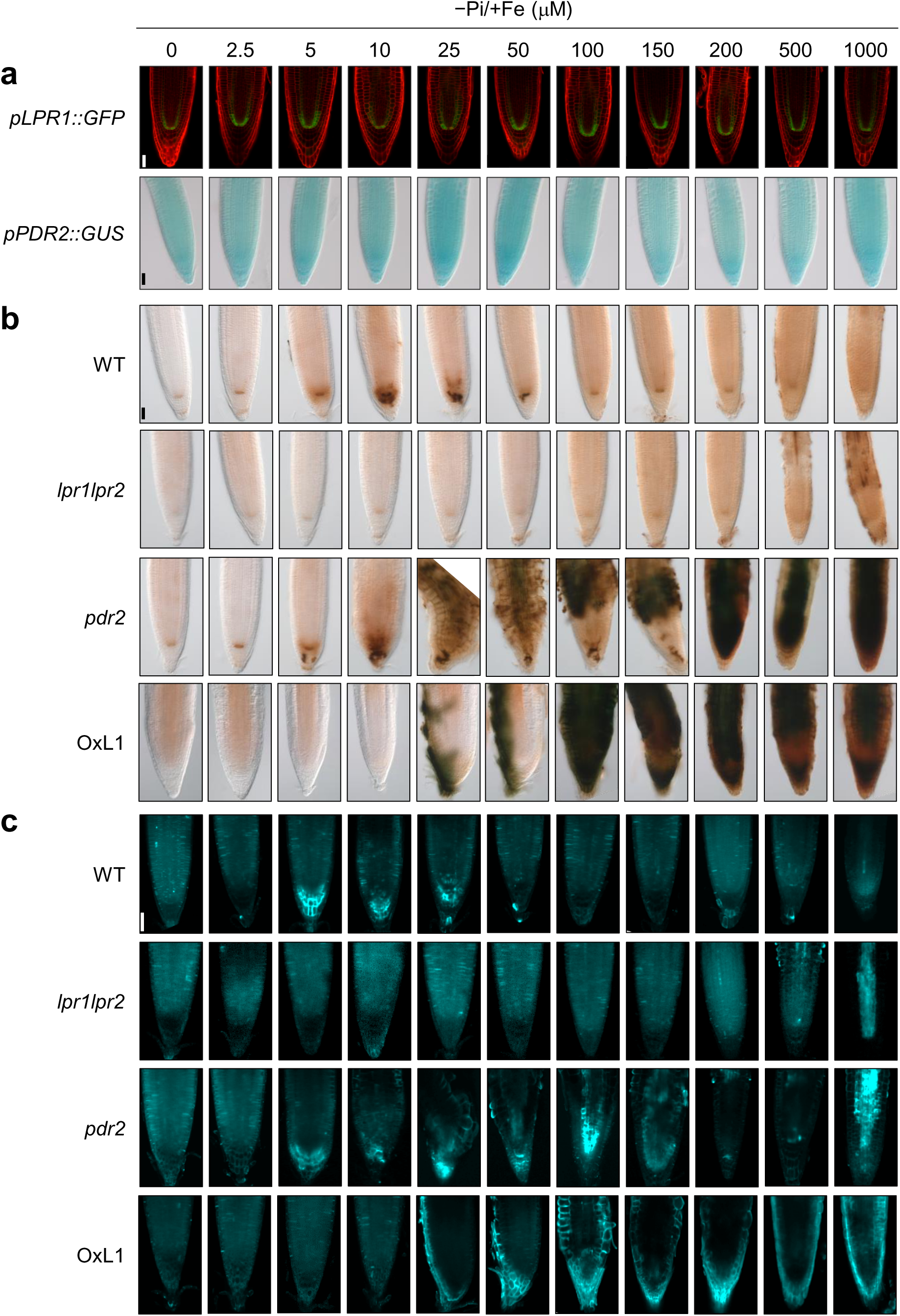
Iron-dependent Fe^3+^ accumulation and callose deposition in Pi-deprived root tips. **a-c.** Seeds of transgenic *pLPR1^Col^::GFP* and *pPDR2^Col^::GUS* lines (wild-type background) as well as of wild-type (WT), *lpr1lpr2, pdr2* and LPR1-overexpression (OxL1) lines were germinated for 5 d on +Pi agar prior to transfer to –Pi media supplemented with increasing iron (Fe^3+^-EDTA) concentrations. After 3 d of transfer, root tips were monitored for *pLPR1::GFP* and *pPDR2::GUS* expression (**a**), for Fe^3+^ accumulation by Perls/DAB staining (**b**), and for callose deposition by aniline blue staining (**c**). Shown are representative images (n≥15). Scale bars 50µm.

### The *PDR2-LPR1* module governs Fe re-distribution in Pi-deprived root meristems

We monitored by Perls/DAB (diaminobenzidine) staining the accumulation and distribution of labile iron (Fe^3+^) in primary root tips upon transfer of Pi-replete seedlings to various –Pi+Fe agar media (Fig. 6b). Tissues of the RAM that accumulate Fe in the apoplast upon Pi deprivation overlap with the cell-specific LPR1 expression domain (Müller et al., 2015). In Pi-starved wild-type roots, Fe progressively accumulated in the SCN with gradually increasing Fe supply. The intensity of Perls/DAB staining peaked at 10 μM Fe and steadily decreased with higher Fe concentrations (25-1000 μM). The RAM of insensitive *lpr1lpr2* seedlings did not stain for iron above background, except for the highest Fe concentrations applied (500 μM and 1000 μM). Interestingly, the cell-type specificity and intensity of Fe staining were evidently similar between Pi-deprived wild-type and *pdr2* root meristems for up to 10 μM Fe supply, while at higher Fe concentrations (25-1000 μM) intense Perls/DAB staining was observed in the entire *pdr2* root tip (Fig. 6b). Importantly, upon transfer of Pi-replete wild-type, *lpr1lpr2*, and *pdr2* seedlings to Fe-supplemented +Pi control media, gradually increasing low external Fe supply (0-10 µM) did not intensify Perls/DAB staining above background (Extended Data Fig. 9a). Moderate Fe supply (25-200 µM) appreciably increased Perls/DAB staining in the SCN and columella of wild-type and *pdr2* root tips, but not in *lpr1lpr2* root meristems, whereas excess Fe supply (500 µM and 1000 μM) caused Fe overload in root tips of all genotypes (Extended Data Fig. 9). Thus, Fe overload and root growth inhibition on excessively high Fe concentration are independent of LPR1 function.

Finally, we monitored callose formation by aniline blue staining in root meristems upon seedling transfer to Fe-supplemented –Pi media. Our data reveal that callose deposition in Pi-deprived root tips largely reflects Fe dose-dependent and genotype-specific patterns of Fe accumulation in root tips (Fig. 6c). The similar responses of Pi-deprived wild-type and *pdr2* root meristems to low external Fe availability (0-10 μM) in terms of growth inhibitions (Fig. 5), Fe^3+^ accumulation (Fig. 6b), and callose deposition (Fig. 6c) are consistent with the conclusion that PDR2 function does not restrict LPR1 expression, biogenesis or ferroxidase activity. However, the unrestrained Fe^3+^ accumulation in *pdr2* root tips at moderately elevated Fe availability (>10 μM) and high external Fe supply suggests a role for PDR2 in maintaining Fe homeostasis and regulating Fe pools in root meristems. This conclusion is supported by the analysis of LPR1-overexpression seedlings (OxL1) for iron-dependent root growth inhibition (Fig. 5d), along with the observed Fe^3+^ accumulation and callose deposition in Pi-deprived root tips (Fig. 6b, c). In both assays, the response of OxL1 roots to increasing external Fe supply mimics the response of *pdr2* roots challenged by Pi limitation. The data suggest that root growth inhibition by gradually increasing low a Fe supply (2.5-25 μM) in –Pi condition is independent of the LPR1 protein level. At intermediate and high Fe concentrations (>25 μM), ectopic *p35S::LPR1* expression shifts Fe^3+^ accumulation from the RAM to the columella and epidermis at agar contact sites, indicating that substrate availability determines LPR1 ferroxidase activity (Fig. 6b).

### Progenitors to embryophytes acquired LPR1-type ferroxidase from soil bacteria

The substantial amino acid sequence similarity (37% identity) between LPR1 and bacterial CotA, which oxidizes bulky organic substrates such as ABTS or bilirubin (Enguita et al., 2004; Sakasegawa et al., 2006), prompted us to study the phylogenetic relationship between LPR1 and annotated MCO proteins (UniProt Database). We retrieved 189 MCO sequences and generated a midpoint-rooted phylogenetic tree featuring two major branches (Fig. 7). MCO group I, composed of two monophyletic clades, assorts fungal laccases including ferroxidases involved in Fe import (clade Ia), and plant laccases with ascorbate oxidases (clade Ib). Paraphyletic MCO group II includes bacterial, fungal, and mammalian MCO proteins of unknown specificities, or of presumed functions in N assimilation (Cu-dependent nitrite reductases), Fe export (ferroxidases), and hemostasis (blood coagulation factors). CotA and LPR1-like MCOs of *Arabidopsis* and rice (Ai et al., 2020) form a monophyletic clade within the bacterial paraphyletic segment of group II (Fig. 7).

**Fig. 7.**
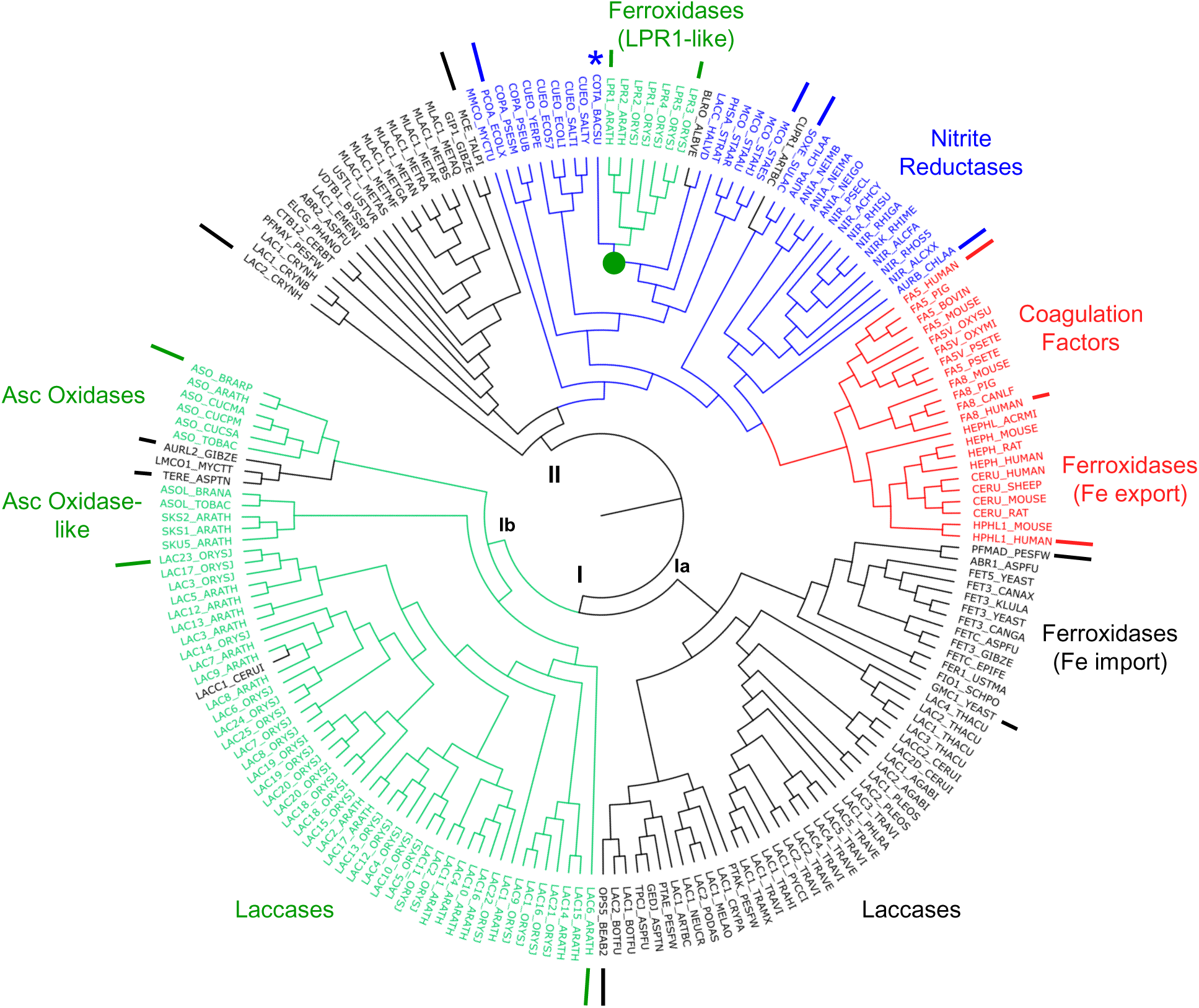
Phylogenetic relationship of MCO proteins. CotA and 187 full-length sequences of annotated MCO proteins (UniProt Knowledge Database) were used to generate a maximum-likelihood midpoint-rooted phylogenetic tree of two major branches. MCO group I includes fungal (clade Ia) and plant (clade Ib) laccases, including ascorbate oxidases and ferroxidases related to Fet3p. Group II includes MCOs of various activities from archaebacteria (black), eubacteria (blue), and animals (red). CotA (*) and LPR1-related ferroxidases form a monophyletic clade within the polyphyletic segment of MCOs from eubacteria.

Comparison of the primary and tertiary structures rationalizes the strikingly different substrate specificities of CotA and LPR1. The amino acid sequence alignment of CotA and LPR1-like MCOs indicates absence of a *bona fide* Fe^2+^-binding acidic triad in CotA (Extended Data Fig. 10). While the first and third acidic amino acid residues on LPR1 (E269 and D462) are embedded in two conserved segments, the second residue (D370) is located in a variable linker flanked by two hydrophobic motifs in LPR proteins (Fig. 3j). Although these features, with the exception of the three acidic residues, are conserved in CotA, its variable linker is shorter (aa 321-326) and likely folds into a tight surface loop, permitting access of bulky molecules (e.g., ABTS) to the substrate binding pocket (Fig. 3d,g, Extended Data Fig. 11a). The longer surface loop on LPR1 (aa 363-373) harbors D370 and may provide a flexible lid-like segment for high-affinity Fe^2+^ binding, in addition to possibly preventing access of bulky substrates (Fig. 3e,h). Notably, despite a similar architecture of the Fe^2+^-binding and electron-transfer sites near the T1 Cu center, the surface topology of the Fe^2+^-binding pocket differs between LPR1 and yeast Fet3p (Extended Data Fig. 11b,c).

Sequence similarity searches (NCBI nucleotide collection) in the eukaryal domain indicated restriction of LPR1-like proteins (presence of the acidic triad) to embryophytes (land plants). Unlike MCO laccases, which form large families in plants (Turlapati et al., 2011), *bona fide* LPR1-like MCO ferroxidases are often encoded by two genes in the bryophytes and tracheophytes (Extended Data Table 2). It is generally accepted that land plants evolved from a small but diverse group of green algae (the charophytes or streptophyte algae), which comprise a paraphyletic assemblage of five classes of mainly freshwater and terrestrial algae (Mesostigmatophyceae including Chlorokybophyceae, Klebsomidiophyceae, Charophyceae, Coleochaetophyceae, and Zygnematophyceae) (de Vries and Archibald, 2018; Furst-Jansen et al., 2020). Phylogenomic analyses increasingly favor the Zygnematophyceae, or alternatively a clade comprising the Zygnematophyceae and Coleochaetophyceae, as the sister group of land plants (Wickett et al., 2014; Zhong et al., 2014). We identified sequences coding for predicted LPR1-type MCOs in the recently published genomes of two Zygnematophyceae (Cheng et al., 2019), *Mesotaenium endlicherianum* and *Spirogloea musicola*, but not in the genomes of *Mesostigma viride* and *Chlorokybus atmophyticus* (Wang et al., 2020) (Extended Data Table 2). We employed a hidden Markov model (HMM) approach using a profile of full-length LPR1-like MCOs from 14 land plants to analyze data sets of the 1KP project (One Thousand Plant Transcriptomes Initiative 2019) (One Thousand Plant Transcriptomes, 2019). We did not identify LPR1-like sequences featuring an acidic triad among the rhodophytes, glaucophytes or chlorophyte algae; however, half of the analyzed charophyte and most of the bryophyte transcriptomes revealed such MCO sequences. Among the charophyte transcriptomes, 23 LPR1-like sequences are present in the Zygnematophyceae and one in the Coleochaetophyceae (Extended Data Table 3, Extended Data Fig. 14).

While the HMM approach supported restriction of LPR1-related proteins to the streptophytes (embryophytes plus charophytes), it pointed to a widespread occurrence of CotA-like MCOs (often annotated as bilirubin oxidases) in the bacterial and archaeal domains (Extended Data Table 3). Interestingly, if filtered for the presence of the LPR1-type acidic triad and variable linker sequence, we identified at least 35 bacterial LPR1-like MCOs with substantial amino acid sequence identity (35-40%) to *Arabidopsis* LPR1 (Extended Data Tables 2-4, Extended Data Fig. 12). We selected four such bacterial MCO sequences for homology modeling, which suggests presence and topology of a LPR1-type surface loop for Fe^2+^ binding (Fig. 3f,i, Extended Data Fig. 13a). Indeed, when expressed in *Escherichia coli*, recombinant LPR1-like MCOs from *Streptomyces clavuligerus* and *Sulfurifustis variabilis* show ferroxidase activity (Extended Data Fig. 13b-c).

Bacterial genera harboring the >35 LPR1-like ferroxidase genes are limited to five phyla (Extended Data Table 2), comprising so-called Terrabacteria (Firmicutes, Actinobacteria, Chloroflexi) as well as members of Bacteroidetes and Proteobacteria isolated from soil habitats (Battistuzzi et al., 2004; Battistuzzi and Hedges, 2009; Marin et al., 2017). A phylogenetic tree of DNA sequences coding for LPR1-like MCOs in bacteria, two streptophyte algae and selected embryphytes reveals a monophyletic clade of streptophyte sequences nested within the radiation of bacterial LPR1-like MCOs (Fig. 8a). Such a tree topology suggests a single horizontal gene transfer (HGT) event from a bacterial donor to a progenitor of the embryophytes, which is consistent with the exon-intron structure of *LPR1*-like genes (Fig. 8b). The number of introns increased from one (bryophytes) to three (tracheophytes) during evolution. All introns are in phase-0 and thought to partition the gene of bacterial origin into symmetric exons for maintaining its ancient functionality (Mayer et al., 2011; Husnik and McCutcheon, 2018). A cladogram including coding sequences of additional bryophytes and streptophyte algae (1 KP Project), which correspond to the internal polypeptide covering the entire acidic triad segment in *Arabidopsis* LPR1 (aa 264-465), supports the proposition that streptophyte ancestors of the embryophytes acquired LPR1-type ferroxidase from soil bacteria by HGT (Extended Data Fig. 14).

**Fig. 8.**
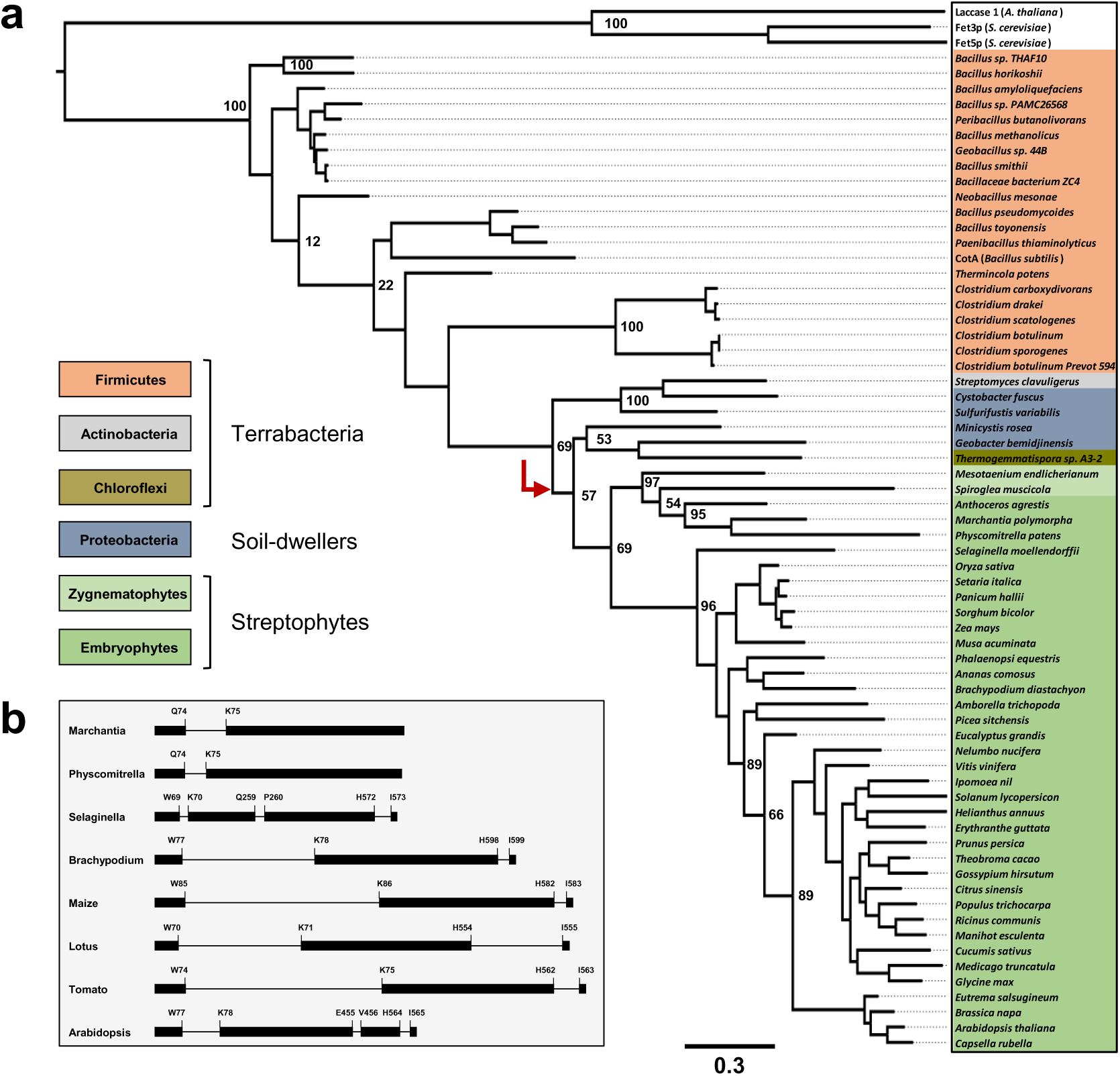
Phylogenetic relationship of LPR1-like MCO genes. **a.** Phylogenetic tree of select LPR1-like MCO coding DNA sequences (CDS) from bacteria, Zygnematophyceae and embryophytes (see Extended Data Table 2). Full-length bacterial CDS and streptophyte CDS starting from the predicted second exon (see panel **b**) were used to generate the maximum-likelihood midpoint-rooted tree (400 bootstrap replicates). The streptophyte sequences occupy a monophyletic clade nested within the paraphyletic bacterial radiation, which suggests a single horizontal gene transfer event from a bacterial donor (red arrow). **b.** The gene models of select LPR1-like genes suggest acquisition of phase-0 introns (separating symmetric exons) to maintain original bacterial gene function.

### LPR1-type MCO ferroxidases emerged during bacterial land colonization

To explore the origin of LPR1-type ferroxidases, we searched, using the internal 202-aa LPR1 segment as query, the eubacterial and archaeal domains for putative MCO sequences harboring only partial acidic triad motifs. We additionally identified at least 80 such MCO sequences, which are absent in marine Hydrobacteria but limited to Terrabacteria, Proteobacteria, and Halobacteria (Extended Data Table 3). Most of the predicted MCOs (∼60) presumably lack the second conserved acidic residue (corresponding to D370 on LPR1) in their variable linker sequences, which however is expendable for high-affinity Fe^3+^-binding (Fig. 4). Predicted MCOs of the Firmicutes show the highest combinatorial variation of acidic triad motifs, suggesting that CotA-type laccases evolved from LPR1-type MCOs by linker contraction and sequential loss of acidic triad residues (Extended Data Table 4). A cladogram comprising bacterial MCO sequences related to LPR1 and CotA suggests that LPR1-type ferroxidases arose early during land colonization and were subject to lateral gene transfer among phyla of Bacteria and Archaea (Extended Data Fig. 15). Because many of the extant bacterial genera identified are known to conduct iron- or sulfur-based anoxygenic photosynthesis and/or to recycle its organic products by dissimilatory iron or sulfate reduction under anaerobic or microaerophilic conditions, LPR1-type MCOs with presumed functions in iron metabolism likely emerged prior to oxygenic photosynthesis by cyanobacteria and the ensuing Great Oxygenation Event (Weber et al., 2006; Sleep and Bird, 2008).

## Discussion

Uneven Pi availability guides root development via local adjustment of root tip growth. Upon Pi limitation, root meristem maintenance is under genetic control of the *PDR2-LPR1* module and depends on Fe co-occurrence (Svistoonoff et al., 2007; Ticconi et al., 2009; Müller et al., 2015). Here we show that *LPR1*, a principal determinant of local Pi sensing in *A. thaliana*, encodes a novel prototypical MCO ferroxidase of high substrate specificity (Fe^2+^) and affinity (*K*_m_ ∼ 2 μM). Metal oxidases including ferroxidases related to Fet3p (limited to Fungi) or ceruloplasmin (limited to Animalia) form a major group within the ancient MCO superfamily whose diverse members are widely distributed in all domains of life (Janusz et al., 2020). Although LPR1 and Fet3p share similar catalytic parameters as well as an analogous architecture of the Fe^2+^-binding and adjoining T1 Cu site (Jones et al., 2020), our study suggests that LPR1-type MCOs displaying ferroxidase activity emerged very early during bacterial land colonization. LPR1-type ferroxidases (or their MCO progenitors) possibly crossed bacterial phyla multiple times to diversify by lateral gene transfer, which is widespread among soil bacteria (Ochman et al., 2000; Klumper et al., 2015). For example, in the genus *Bacillus* (Firmicutes), LPR1-type MCOs evolved to CotA-type laccases by progressive remodeling of the acidic triad segment. Gram-positive soil bacteria produce endospores, which are fortified with various spore coat proteins including CotA to survive in harsh environments (McKenney et al., 2013). While the precise biochemical function of CotA is not known, its unusually large substrate binding cavity and correspondingly short lid-like loop, which likely derived from an LPR1-type progenitor MCO (Extended Data Table 3, Extended Data Fig. 15), presents a unique structural feature among MCO laccases (Enguita et al., 2003).

The physical proximity between soil bacteria and the terrestrial/subaerial common ancestor of streptophytes likely facilitated our proposed single HGT event of LPR1-type ferroxidases, which occurred before the divergence of Zygnomatophyceae (possibly extending to Coleochaetophyceae) and embryophytes (∼580 mya) (Cheng et al., 2019). Our data corroborate the conjecture that plant terrestrialization was accelerated by substantial HGT from soil bacteria to early land plant progenitors (Yue et al., 2012; Husnik and McCutcheon, 2018; Cheng et al., 2019). Diversification of LPR1-type ferroxidases in land plants was possibly limited to rare whole-genome duplication events (Soltis and Soltis, 2016), which is suggested by the single retained *LPR* sister gene pair in *A. thaliana*, *LPR1* (At1g23010) and *LPR2* (At1g71040) (Abel et al., 2005). The unknown extent of lateral gene transfer among bacteria curtails precise identification of an HGT donor, probably explaining why LPR1-type ferroxidase sequences of four bacterial phyla (Proteobacteria, Chloroflexi, Actinobacteria, Firmicutes) are monophyletic with the streptophyte sequences (Fig. 8a, Extended Data Fig. 15). Nonetheless, the metabolic lifestyle and iron biochemistry of extant bacterial sister genera may allow insight into the function of plant LPR1-like ferroxidases. Members of the four phyla are facultative anaerobic or microaerophilic, spore-forming chemoorganotrophs, which are capable of dissimilatory or fermentative Fe^3+^ reduction and have been isolated from iron-rich soils or artificial Fe(III) oxide-enriched growth substrates (Lentini et al., 2012; Li et al., 2012; List et al., 2019; Wang et al., 2020). For example, genera of the Geobacteraceae family, including *Geobacter* and closely related *Desulfuromonas* species, are the predominant Fe^3+^ reducers in many anaerobic sediments. Such bacteria chemotactically locate extracellular Fe(III) oxide minerals and transfer electrons via nanowires to its solid-phase surfaces (Childers et al., 2002). Other microbial strategies for accessing insoluble Fe(III) oxides involve production of soluble external electron shuttles such as redox-active antibiotics or chelating ligands, which simultaneously increase Pi bioavailability in soil (Reguera et al., 2005; Weber et al., 2006; Liptzin and Silver, 2009; Glasser et al., 2017; Michelson et al., 2017; McRose and Newman, 2021) (Fig. 9). Although considered as strict anaerobes, *Geobacter* species can tolerate episodes of dioxygen exposure and code for ROS-scavenging proteins (Methe et al., 2003). Notably, *G. metallireducens* contains four genes encoding CotA-like proteins, the genes presumably having been acquired from *Bacillus* by lateral gene transfer (Berini et al., 2018). While two of these genes encode LPR1-type MCOs (see Extended Data Fig. 15), a fifth gene expresses a biochemically characterized ABTS-oxidizing MCO with a very low *K*_m_ for dioxygen (<10 M) (Berini et al., 2018). If low *K*_m_ (O_2_) values also apply to similar MCO enzymes, bacterial LPR1-type ferroxidases may promote Fe-redox cycling to protect against oxidative stress associated with Fe^3+^ reduction and resultant Fenton chemistry.

**Fig. 9.**
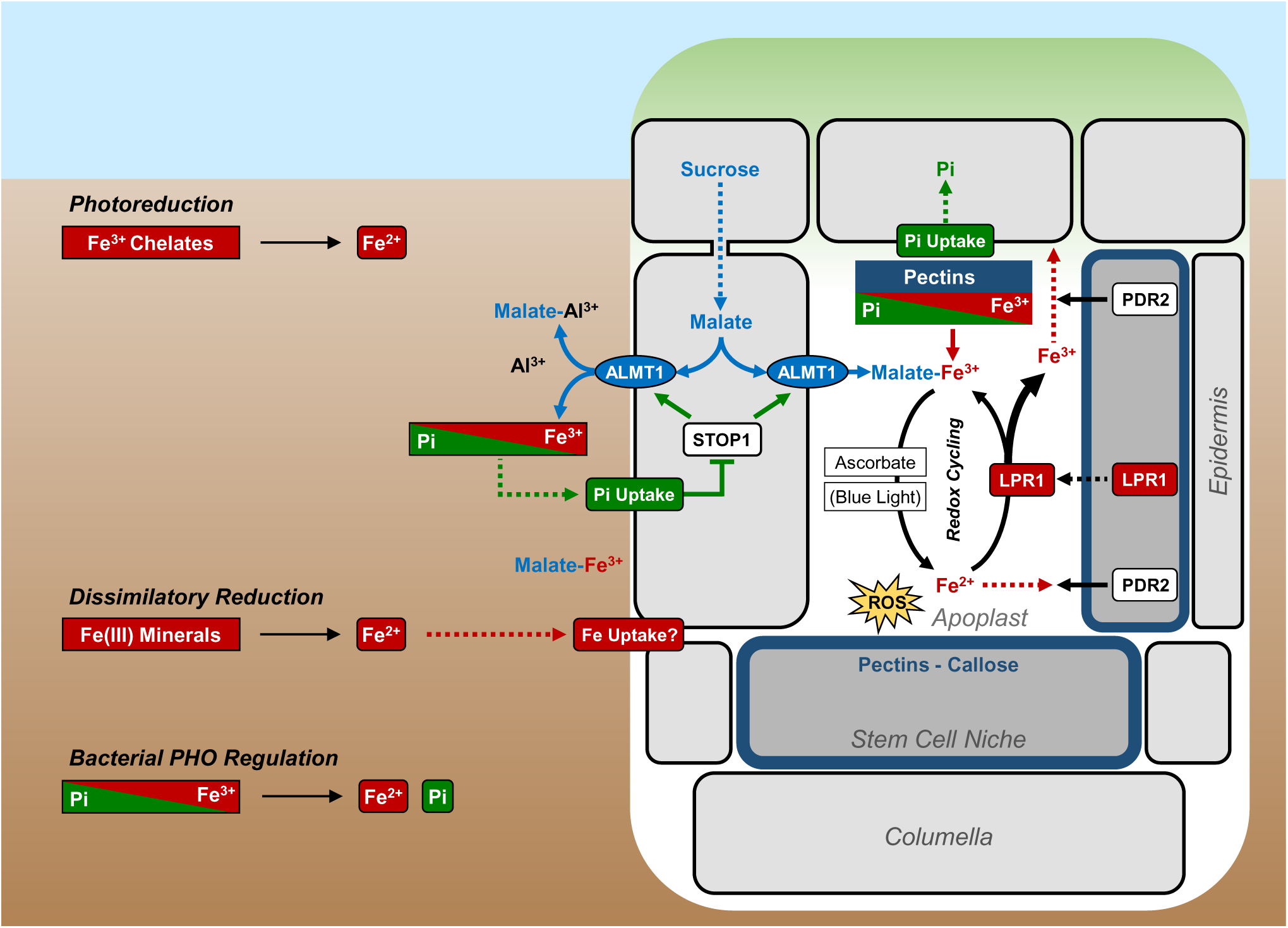
Model of LPR1 function in the context of microbial soil chemistry. The STOP1-ALMT1 module activates on low Pi the release of malate into the rhizosphere and apoplast of internal root tissues, where malate mobilizes Pi from pectin-associated Fe-Pi complexes, and chelates toxic Al^3+^ cations in soil (Kochian et al., 2015; Balzergue et al., 2017; Mora-Macias et al., 2017). Ascorbate readily reduces Fe^3+^-malate species in cell walls (Grillet et al., 2014), stimulating ROS formation by Fenton chemistry. LPR1-dependent Fe^2+^ oxidation and Fe redox cycling attenuate ROS production and presumably ROS signaling in the stem cell niche (SCN). Fe^2+^ substrate availability tunes the ferroxidase activity of the constitutively expressed LPR1 protein. PDR2, the single orphan, ER-resident P5-type ATPase in *Arabidopsis* (AtP5A) counteracts LPR1 function by maintaining Fe homeostasis in root tips (e.g., by promoting unknown apoplastic and/or symplastic Fe export processes). We propose that Fe^2+^ generation in the SCN apoplast, and possibly Fe^2+^ uptake from the rhizosphere into the columella/SCN apoplast, constitute a local cue for external Pi availability monitored by the PDR2-LPR1 module. Blue light-dependent Fe^3+^ photoreduction produces ROS and inhibits root tip growth in low Pi under controlled laboratory conditions (i.e. growth on sterile, light-exposed agar plates) (Zheng et al., 2019). However, in such settings, Fe^3+^ photoreduction likely mimics the impact of bacterial communities in the rhizosphere, which mobilize redox-active Fe^2+^ from Fe(III) oxide minerals by various metabolic processes. These include dissimilatory Fe^3+^ reduction (facilitated by ligand chelation, extracellular electron shuttles, or nanowire formation), and *PHO* regulon-dependent Pi desorption from Fe(III) minerals via Fe^3+^ reduction by extracellular, redox-active antibiotics (Weber et al., 2006; Glasser et al., 2017; McRose and Newman, 2021).

LPR1 and related plant ferroxidases likely facilitate analogous processes in root tips. Upon Pi limitation, the STOP1-ALMT1 module, a C_2_H_2_ zinc finger transcription factor (SENSITIVE TO PROTON RHIZOTOXICITY 1) and one of its direct target genes (*ALUMINUM-ACTIVATED MALATE TRANSPORTER 1*), activates malate release into the rhizosphere and apoplast of internal root tissues (Balzergue et al., 2017) (Fig. 9). Malate mobilizes Pi from insoluble metal complexes by Fe^3+^ chelation (Abel, 2017; Balzergue et al., 2017; Mora-Macias et al., 2017; Gutierrez-Alanis et al., 2018). Ensuing cell wall chemistry (e.g., Fe^3+^ reduction by ascorbate), augmented by photochemistry under laboratory conditions, generate Fe^2+^ and ROS (Grillet et al., 2014; Müller et al., 2015; Abel, 2017; Zheng et al., 2019). LPR1-dependent Fe redox cycling (Fe^3+^ re-formation and cell wall deposition) attenuates ROS production and presumably ROS signaling in the SCN. We propose that the constitutively expressed MCO ferroxidase, LPR1, senses subtle increases in Fe availability as a Pi-dependent cue to adjust root tip growth to Pi deprivation. Regulation of LPR1 activity by substrate availability is supported by our observation that PDR2/AtP5A counteracts LPR1 function by maintaining Fe homeostasis in root meristems (Fig. 9), which points to a novel role of single, ER-resident, orphan P5-type ATPases in plants (Sorensen et al., 2015; Naumann et al., 2019).

## Methods

### Plant lines and growth conditions

*Arabidopsis thaliana* accession Columbia (Col-0), Col-0 mutant lines *pdr2-1*, *lpr1lpr2*, and transgenic lines *pCaMV 35S::LPR1 (p35S::LPR1)* and *pLPR1::eGFP-GUS* were previously described (Svistoonoff et al., 2007; Ticconi et al., 2009; Müller et al., 2015). GATEWAY technology (Invitrogen) and *Agrobacterium*-mediated transformation were used to generate transgenic *Arabidopsis* lines expressing *p35S::LPR1^E269A^; p35S::LPR1^D370A^* and *p35S::LPR1^D462A^*. Seeds were surface-sterilized and germinated on 1% (w/v) Phyto-Agar (Duchefa) containing 2.5 mM KH_2_PO_4_, pH 5.6 (high Pi or +Pi medium) or no Pi supplement (low Pi or –Pi medium), 50 µM Fe^3+^-EDTA, 5 mM KNO_3_, 2 mM MgSO_4_, 2 mM Ca(NO_3_)_2_, 2.5 mM MES-KOH, pH 5.6, 70 μM H_3_BO_3_, 14 μM MnCl_2_, 10 μM NaCl, 0.5 µM CuSO_4_, 1 µM ZnSO_4_, 0.2 µM Na_2_MoO_4_, 0.01 µM CoCl_2_ and 5 g/l sucrose. The agar was routinely purified by repeated washing in deionized water and subsequent dialysis using DOWEX G-55 anion exchanger (Ticconi et al., 2009). ICP-MS analysis of the washed agar (7.3 μg/g Fe and 5.9 μg/g P) indicated a contribution of 1.3 μM Fe and 1.9 μM P to the solid 1% agar medium. For root length measurements, 27-54 seedlings were transferred to the indicated media and gain of primary root length was marked daily. Photos were analyzed using ImageJ software. Additional lateral root were induced as previously described (Himanen et al., 2002). Hydroponically grown seedlings were germinated under moderate shaking in 200-ml flasks containing 50 ml liquid +Pi medium.

### Microscopy

Green fluorescent protein (GFP) fluorescence was visualized using a Zeiss LSM 780 confocal laser-scanning microscope (excitation 488 nm, emission 536 nm) in phosphate-buffered saline. Co-localization of GFP and PI (propidium iodide) was monitored in sequential mode (excitation 561 nm, emission 630 nm). Seedlings were incubated for 2 min in 0.1 mg/ml PI solution. For GUS (β-glucuronidase) staining, seedlings were incubated in 50 mM Na-phosphate (pH 7.2), 0.5 mM K_3_Fe(CN)_6_, 0.5 mM K_4_Fe(CN)_6_, 2 mM X-Gluc, 10 mM EDTA and 0,1% TritonX at 37°C and subsequently cleared using chloral hydrate solution (7:7:1 chloral hydrate:ddH_2_O:glycerol) as described before (Wong et al., 1996). Callose was stained for 1 h with 0.1% (w/v) aniline blue (AppliChem) in 100 mM Na-phosphate buffer (pH 7.2) and carefully washed twice. Fluorescence was visualized using a Zeiss LSM 880 confocal laser-scanning microscope (excitation 405 nm, emission 498 nm) in 100 mM Na-phosphate buffer (pH 7.2)(Müller et al., 2015). Histochemical iron staining (Perls/DAB) was performed as previously described (Müller et al., 2015) with minor changes to the protocol. Plants were incubated for 10 min in 4% (v/v) HCl, 4% (w/v) K-ferrocyanide (Perls staining), or K-ferricyanide (Turnbull staining). For DAB intensification, plants were washed twice (ddH_2_O) and incubated (15 min) in methanol containing 10 mM Na-azide and 0.3% (v/v) H_2_O_2_. After washing with 100 mM Na-phosphate buffer (pH 7.4), plants were incubated for 3 min in the same buffer containing 0.025% (w/v) DAB (Sigma-Aldrich) and 0.005% (v/v) H_2_O_2_. The reaction was stopped by washing 100 mM Na-phosphate buffer (pH 7.4) and optically clearing with chloral hydrate (1 g/ml, 15% glycerol).

### Real-time quantitative PCR

Total RNA was prepared from excised root tips (tip growth gained after seedling transfer) by using the peqGOLD Plant RNA Kit (VWR). One biological replicate represents 40-60 pooled root tips. RNA samples were dsDNase treated (dsDNase, Thermo Scientific, EN0771) and quantified. cDNA was prepared using 1 µg total RNA, which was reverse transcribed by using oligo(dT) with a First-Strand cDNA Synthesis Kit (Fermentas) according to the manufacturer’s protocol. Quantitative real-time PCR was performed with first strand cDNA template on a QuantStudio 5 PCR System with Fast SYBR Green Mix (Applied Biosystems). The reported fold induction was analyzed by the ΔCt-method and normalized to the endogenous UBC9 (ubiquitin-conjugating enzyme 9) control. Gene-specific amplimers are listed in Supplementary Table 1.

### Quantitative proteomics

Primary root tips were excised and collected into liquid nitrogen. Tissue lysis, sample preparation, protein labeling, Tandem-Mass-Tag (TMT) spectrometry, MS/MS data analysis, and TMT-quantifications were performed as recently described (Rodriguez et al., 2020; Stephani et al., 2020).

### LPR1 homology modeling

YASARA 13.9 (Krieger et al., 2002; Krieger et al., 2009) was used to derive 25 homology models of LPR1 or bacterial MCOs, each based on five PDB (The Protein Data Bank) (Berman et al., 2000) templates (five X-ray structures of laccase CotA from *B. subtilis*: 2WSD, 2X88, 4AKO, 2X87, and 4AKP). Quality analysis with PROCHECK (Laskowski, 1993) and PROSA II (Sippl, 1990, 1993) identified the best fit for each protein. All Cu^+^ cations of the templates were adopted and merged into the models. The ferrous iron (Fe^2+^) was manually added in proximity to residues E269 and D370 of the highly conserved acidic triad on LPR1 or the respective conserved positions in the bacterial ferroxidase models. Subsequently, the model was refined by 20 cycles of simulated annealing refinement with the corresponding tool of YASARA. Molecular surfaces were created with the modeling program MOE (Molecular Operating Environment v2019.0101, Chemical Computing Group Inc., Montreal, QC, Canada, 2019).

### Site-directed mutagenesis

For introducing point mutations into plasmid DNAs, site-directed mutagenesis was carried out with the Quick Change II Site-directed mutagenesis Kit (Agilent) according to the manufacturer’s instructions. Briefly, two complementary primers containing the desired mutation of the plasmid were used to amplify two overlapping, complementary strands of the plasmid with staggered nicks. After amplification, the parental DNA was digested with *Dpn* I and the mutated plasmids were transformed into *E. coli* Top 10 or XL1 Blue cells. The primers used for generating the different mutations are listed in Supplementary Table 1.

### Purification of native LPR1 protein variants

Transgenic *A. thaliana lpr1* mutant plants expressing *p35S::LPR1*, *p35S::LPR1^E269A^; p35S::LPR1^D370A^* and *p35S::LPR1^D462A^* were grown for 8 weeks on soil in short-day conditions (8 h light, 16 h darkness, 21°C). Entire plant rosettes were harvested and homogenized in liquid nitrogen. To extract whole proteins, 15 g plant material was vortexed in 40 ml buffer A (50 mM Tris-Cl pH 6.8, 100 mM NaCl, 0.5 mM EDTA, 10% glycerol) containing 1 mM PMSF and 1× protease inhibitor (ROCHE) followed by incubation for 30 min at 4°C (shaking). After clearance of the extract by centrifugation (500 × *g*, 30 min, 4°C), the supernatant was subjected to 40% (NH_4_)_2_SO_4_ precipitation (1 h at 4°C). The resulting pellet (4,500 × *g*, 45 min, 4°C) was discarded and the supernatant treated with 80% (NH_4_)_2_SO_4_ for 1 h at 4°C. The resulting protein pellet was solubilized in 2-3 ml buffer A, loaded on a HighLoad Superdex 200 gel filtration column (HL 16/60, GE Healthcare), and eluted with buffer A as the mobile phase. Fractions containing LPR1 (detected by immune blot analysis) were directly applied to cation exchange carboxymethyl-sepharose column (HiTrap CM FF, 1-ml, GE Healthcare) equilibrated with buffer B (20 mM Na_2_HPO_4_-NaH_2_PO_4_, pH 7). Fractions were eluted using a linear salt gradient (0-1 M NaCl) in buffer B. LPR1 eluted at 350 mM NaCl. LPR1-containing fractions were stored at −20°C until further use and the enzyme was stable for two weeks. LPR1 abundance and activity were confirmed by immunoblot analysis and ferroxidase assays, respectively. For protein silver staining, the gels were incubated twice for 20 min each or overnight in fixing solution (10% acetic acid, 40% methanol). Subsequently, the gels were incubated in 30% methanol, 1.2 mM NaS_2_O_3_, 829 mM Na-acetate for 30 min, followed by three washing steps (5 min each) in distilled H_2_O. Silver staining was performed by incubating the gels in an aqueous AgNO_3_ solution (2 mg/ml) for 20 min followed by two washing steps with water. The gels were developed (staining of protein bands) in 236 mM NaCO_3_ containing 0.04% formaldehyde, and the reaction was stopped by incubation in 40 mM Na-EDTA.

### Deglycosylation and phosphatase treatments

Purified LPR1 protein was analyzed using Protein Deglycosylation Mix II (New England Biolabs) according to the manufacturer’s instructions. Fetuin was used as a control. Phosphorylation of purified LPR1 protein was tested according to (Maldonado-Bonilla et al., 2014) using λ-protein phosphatase (New England Biolabs). In brief, root material was harvested in phosphatase buffer supplemented with 1 × protease inhibitor (ROCHE). After addition of 1,200 U phosphatase, reactions were carried out for 90 min at room temperature. Samples were inactivated at 95°C for 5 min and analyzed by immunoblotting.

### Peptide sequencing

Proteins were in-gel digested with trypsin and further processed as previously described (Majovsky et al., 2014). Dried peptides were dissolved (5% acetonitrile/0.1% trifluoric acid), injected into an EASY-nLC 1000 liquid chromatography system (Thermo Fisher Scientific), and separated by reverse-phase (C18) chromatography. Eluted peptides were electro-sprayed on-line into a QExactive Plus mass spectrometer (Thermo Fisher Scientific). A full MS survey scan was carried out with chromatographic peak width. MS/MS peptide sequencing was performed using a Top10 DDA scan strategy with HCD fragmentation. MS scans with mass to charge ratios (m/z) between 400 and 1300 and MS/MS scans were acquired. Peptides and proteins were identified using the Mascot software v2.5.0 (Matrix Science) linked to Proteome Discoverer v 2.1 (Thermo Fisher Scientific). A precursor ion mass error of 5 ppm and a fragment ion mass error of 0.02 Da were tolerated in searches of the TAIR10 database amended with common contaminants.

Carbamidomethylation of cysteine was set as fixed modification and oxidation of methionine was tolerated as a variable modification. A spectrum (PSM), peptide and protein level false discovery rate (FDR) was calculated for all annotated PSMs, peptide groups and proteins based on the target-decoy database model and the percolator module. PSMs, peptide groups and proteins with q-values beneath the significance threshold α=0.01 were considered identified.

### Immunoblot analysis

Polyclonal LPR1 epitope-specific antibodies were raised in rabbits against a mixture of two synthetic peptides (peptide I: 175-PKWTKTTLHYENKQQ-189; peptide II: 222-VESPFQLPTGDEF-234) and affinity-purified (EUROGENTEC, Seraing, Belgium). Total proteins were extracted from frozen plant material in buffer A (50 mM Tris-HCl, pH 6.8, 100 mM NaCl, 0.5 mM EDTA, 10% glycerol) containing 1 × protease inhibitor (ROCHE). After centrifugation (20,000 × *g*, 10 min, 4 °C), the protein concentration of the supernatant was determined (2D-Quant, GE Healthcare), and proteins were separated by SDS/PAGE on 8-10% (w/v polyacrylamide) gels and transferred to PVDF membranes (Semi-Dry-Blot, GE Healthcare). After transfer, membranes were exposed to blocking buffer (1× TBS, 0.05% w/v Tween, 3% w/v milk powder) at room temperature for 1 h or overnight. To detect LPR1, affinity-purified, peptide-specific anti-LPR1 antibody was used 1:1000 in blocking buffer for 1 h at room temperature or at 4 °C overnight. Horseradish-peroxidase-conjugated goat anti-rabbit IgG (BioRad, 1:5000) was chosen as a secondary antibody, and the ECL Select or Prime Western Blotting Detection Reagent (Thermo Fisher) was used for visualization. The epitope-specific anti-LPR1 antibody detects 100 ng purified, native LPR1 protein and recognizes only one major protein of ca. 70 kDa in extracts of the *p35S::LPR1* overexpression line (Extended Data Fig. 2). Plant specific actin-antibody (Sigma-Aldrich) was used (at a dilution of 1:2000) as loading control. To control for recombinant bacterial MCO expression anti-His-HRP (Miltenyi Biotec) was used (at a dilution of 1:10000) in the blocking buffer.

### Ferroxidase and other MCO assays

Protein concentration was determined using Qubit Fluorometric Quantification System (Thermo Fisher) according to the manufacturer’s instructions. All reagents except human ceruloplasmin (Athens Research) were purchased from Sigma-Aldrich. Ferroxidase activity was determined as previously described (Müller et al., 2015) using typically 25 µM Fe(NH_4_)_2_(SO_4_)_2_ × 6H_2_O as the substrate and 3-(2-pyridyl)-5,6-bis(2-[5-furylsulfonic acid])-1,2,4-triazine (ferrozine) as a specific Fe^2+^ chelator to scavenge the remaining substrate after the reactions. The rate of Fe^2+^ oxidation was calculated from the decreased absorbance at 560 nm using a molar extinction coefficient of ε_560_=25,400 M^-1^ cm^-1^ for the Fe^2+^-ferrozine complex(Hoopes and Dean, 2004). Phenol oxidase (laccase) activity with ABTS (2,2’-azino-bis(3-ethylbenzothiazoline-6-sulfonic acid), ascorbate oxidase activity, and bilirubin activity were measured in 0.1 M NaH_2_PO_4_-Na_2_HPO_4_ (pH 5.6 – 7.2) as described (Johannes and Majcherczyk, 2000; Sakasegawa et al., 2006; Peng et al., 2015).

### Transient expression assays

The transient transformation of *Nicotiana benthamiana* leaves was carried out using *Agrobacterium tumefaciens* strains that carried the indicated plasmids and the pCB301-p19 helper plasmid (Burstenbinder et al., 2013). Bacteria were grown overnight to an OD_600_ = 0.5 – 0.8, harvested (10,000 × *g*, 4 min, 4 °C) and washed two times with 2 ml transformation buffer (10 mM MES-KOH, pH 5.5, 10 mM MgCl_2_, 150 µg/ml acetosyringone) and subsequently dissolved in transformation buffer to an OD_600_ of 1. The bacteria carrying the expression construct were mixed 1:1 with the ones harboring the pCB301-p19 plasmid and incubated for 1 h at 20 °C. Subsequently, the bacteria were injected at the bottom side of leafs of 5-7 week-old plants. Samples were harvested 4 d post infiltration.

### Expression of bacterial MCO proteins

Genes encoding potential bacterial ferroxidases from *Sulfurifustis variabilis* (BAU47383.1) and *Streptomyces clavuligerus* (QCS10718.1) were codon optimized (Supplementary Table 2), synthesized at the Invitrogen GeneArt Gene Synthesis platform, and cloned into pVp16-Dest vector for IPTG (isopropyl-β-thiogalactopyranoside)-induced expression (3 h at 37°C) in *E.coli* strain ArcticExpress (Agilent). After sonication, the cleared cell lysates were directly used for ferroxidase activity assays.

### Phylogenetic analyses

For MCO sequence alignment and phylogenetic analysis, 193 referenced protein sequences of the annotated “multicopper oxidase family” were obtained from uniProt Knowledgebase (www.uniprot.org) and filtered for fragments. CotA (P07788) was added to the dataset. All phylogenetic trees were calculated by sequence alignment using MAFFT 7 (Katoh et al., 2002) with default settings and created at the CIPRES web-portal with RAxML 8.2.10 (Stamatakis et al., 2008) for maximum likelihood analyses using the JTT PAM matrix for amino acid substitutions in RAxML.

### Data base searches

Full-length protein sequences related to *Arabidopsis* LPR1 (At1g23010) were obtained for select land plant species from NCBI (tblastn searches). To identify *LPR1*-related genes in the main taxonomic groups of bacteria, archaea, early eukaryotes and basal plants, blastp (version 2.10.1+) (Camacho et al., 2009) and hmmer (version 3.3) (Madera and Gough, 2002) were used (www.hmmer.org) (Camacho et al., 2009; Finn et al., 2011). For bacteria and archaea, representative genomes of the key taxonomic groups were obtained from NCBI assembly. Transcriptome data from the 1KP project (One Thousand Plant Transcriptomes Initiative 2019) (One Thousand Plant Transcriptomes, 2019) were taken for early eukaryotes and basal plant species. The *Arabidopsis* LPR1 protein sequence was used as a query to search each genome or transcriptome. Only hits with an alignment length of >175 amino acids were considered. In addition, a profile hidden Markov models approach was applied, generating an HMM profile for LPR1-releated sequences from 14 higher plant species for scanning each genome or transcriptome. All hits were scanned for the presence of a LPR1-type Fe^2+^-binding site, which is composed of an acidic triad with the following three consensus sequence motifs: 1. [WVI]XP[EA][YAF]X[GA]; 2. N[DTS][AG]XXP[YF]PXG[DE]X(5-10)[VI][ML]XF; and 3. NXTX[DEG]XHP. For final validation, candidate sequences were individually aligned with *Arabidopsis* LPR1 and visually inspected. If applicable, representative contiguous sequences covering all acid triad signature motifs were used as query to interrogate (tblastn searches at NCBI) each bacterial phylum for LPR1-type sequences with incomplete acid triad signatures.

### Statistical Analyses

Statistical differences were assessed by one-way ANOVA and Tukey’s HSD posthoc test, using built-in functions of the statistical environment R (R Development Core Team, 2018). Different letters in graphs denote statistical differences at P < 0.05. Graphs were generated using the ggplot2 R package.

## Acknowledgements

We thank M. Ried for insightful and critical reading of the manuscript, T. Desnos for initial discussions, T. Toev for generating plant lines, G. Durnberger and M. Schutzbier, Vienna BioCenter, for expert technical assistance, and members of the department for suggestions. This work was supported by institutional core funding (Leibniz Association) from the Federal Republic of Germany and the State of Saxony-Anhalt to S.A. and by an EMBO short-term fellowship to C.N. Work in the K.M. group was financially supported by the EPIC-XS, project number 823839, the Horizon 2020 Program of the European Union, and the ERA-CAPS I 3686 project of the Austrian Science Fund.

## Author contributions

C.N. and S.A. conceived and designed the experiments; C.N. and M.H. conducted the major experiments and analyzed the data; C.A., N.T., A.T.N., J.Z. and S.A. contributed to additional experiments or data analysis; W.B. performed the homology modeling; R.I., K.M., Y.D., and W.H. conducted and advised the proteomics analyses; P.J. and M.Q. conducted and advised the phylogenomic analyses; G.S. provided conceptual insight and edited the article; C.N. and S.A. wrote the article.

---

**Extended Data Fig. 1.**
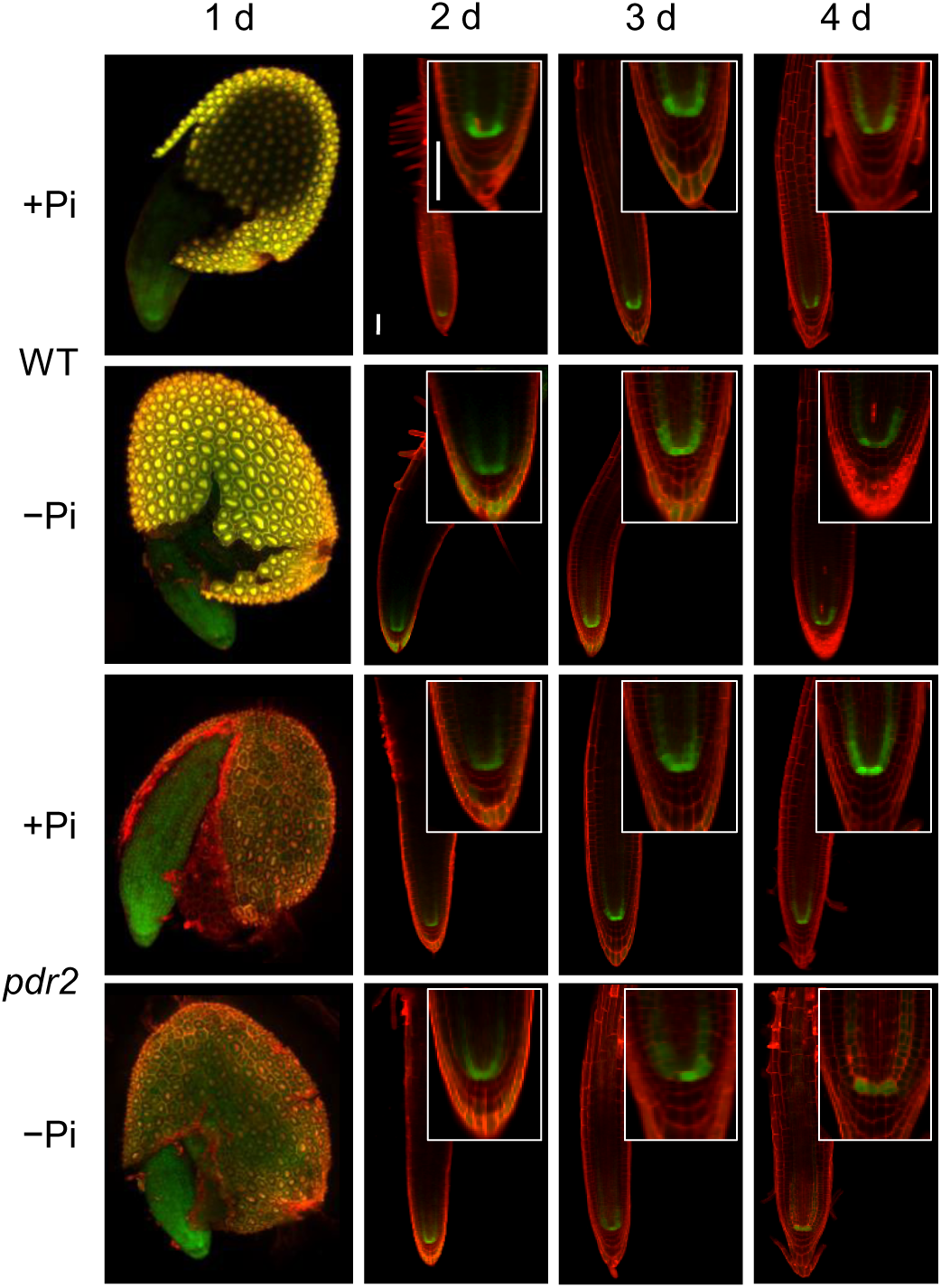
LPR1 expression in root meristems is independent of PDR2 and Pi availability. Expression of *pLPR1^Col^::GFP* in primary root tips of transgenic wild-type (WT) and *pdr2-1* (*pdr2*) plants. Seeds were continuously germinated on +Pi or –Pi agar medium for up to 4 d. Roots were counterstained with PI (red fluorescence), and GFP-derived fluorescence (green) was analyzed. Shown are representative images (n≥10). Scale bars, 50 µm. Whole seed images were generated by Z-Stack fusion.

**Extended Data Fig. 2.**
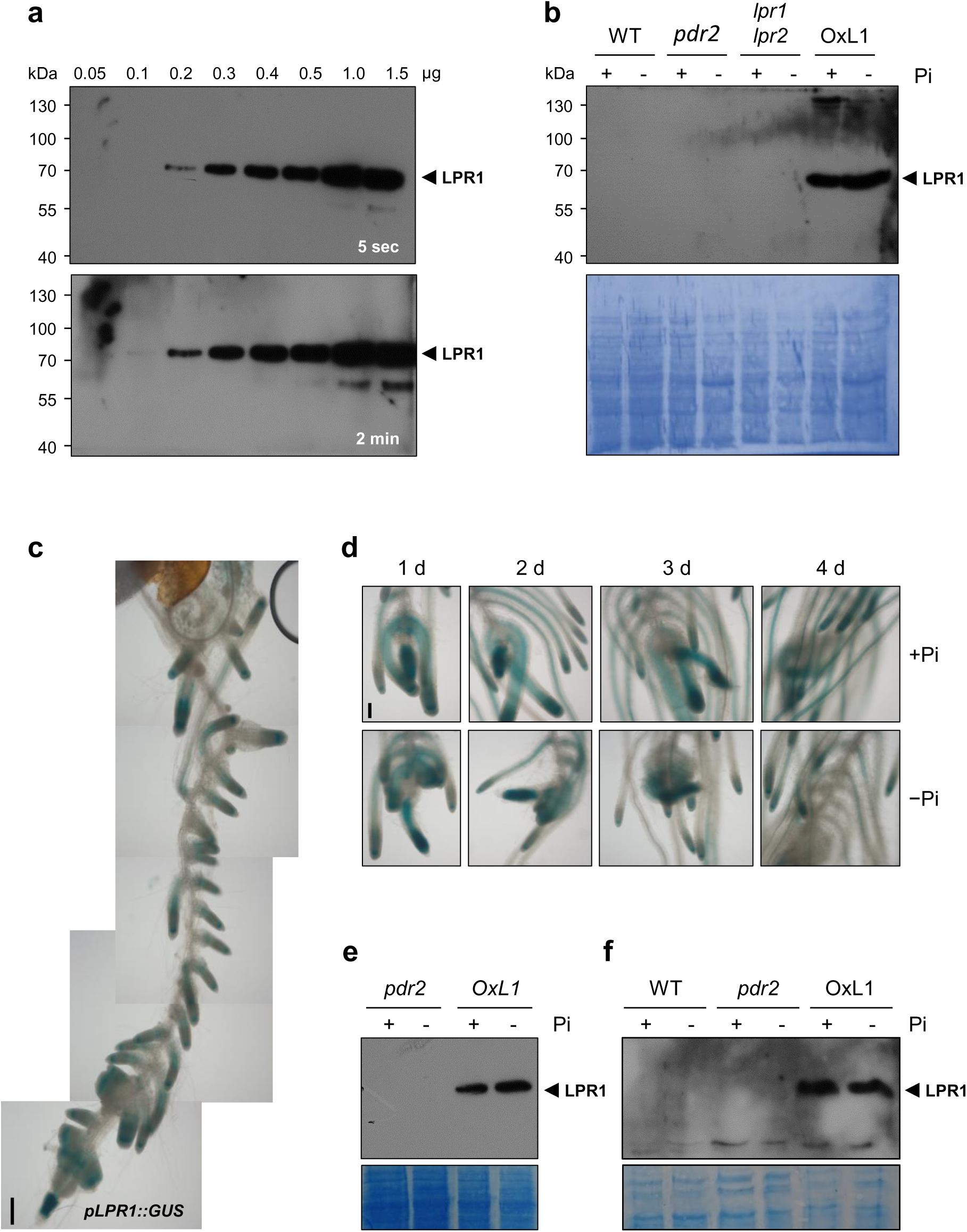
Specificity and sensitivity of epitope-specific anti-LPR1 antibodies. **a.** Increasing amounts of purified, native LPR1 protein were separated by SDS-PAGE on 8% gels, transferred to a PVDF membrane and probed by immunoblot analysis. **b.** Immunoblot analysis of LPR1 expression in Pi-replete (+Pi) and Pi-deprived (–Pi) root extracts of wild-type (WT), *pdr2*, *lpr1lpr2*, and transgenic *p35S::LPR1* seedlings (OxL1). Seeds were germinated on +Pi agar medium (5 d) and transferred to +Pi or –Pi medium for 1 d. Root tips were harvested and root extracts (104 μg total protein) were blotted and analyzed (anti-LPR1 antibodies). Membranes were stained with Coomassie-Blue (n=3). **c.** Expression of *pLPR1^Col^::GUS* in lateral root tips of transgenic wild-type plants stimulated by an auxin-based induction system using NPA-NAA. Seeds were germinated on +Pi medium supplemented with 10 µM NPA for 3 d, prior to transfer to +Pi supplemented with 10 µM NAA to induce lateral root development. After 4 d of transfer, root tips were monitored for GUS expression. Shown is one composite representative image. Scale bars, 100 µm. **d.** Expression of *pLPR1^Col^::GUS* in lateral root tips of transgenic wild-type plants stimulated by an auxin-based induction system using NPA-NAA in response to Pi. Seeds were germinated for 3 d on +Pi medium supplemented with 10 µM NPA, prior to transfer to +Pi supplemented with 10 µM NAA to induce lateral root development. After 4 d of transfer, seedlings were transferred for up to 4 d to +Pi or –Pi medium without supplements. Root tips were monitored for GUS expression. Shown are representative images (n≥10). Scale bars, 100 µm. **e.** Immunoblot analysis of LPR1 expression in root extracts prepared from Pi-replete (+Pi) and Pi-deprived (–Pi) seedlings of *pdr2* and transgenic *p35S::LPR1* plants (OxL1). Seeds were germinated in +Pi liquid medium for 7 d prior to the addition of 10 µM NPA for 3 d. After media exchange, seedlings were treated with 10 µM NAA for 6 d, washed and transferred to liquid +Pi or –Pi media without supplements for 3 d. Whole root extracts (51 μg total protein) were blotted and analyzed (anti-LPR1 antibodies). Membranes were stained with Coomassie-Blue (n=4). **f.** Immunoblot analysis of LPR1 expression in pre-fractionated root extracts prepared from Pi-replete (+Pi) or Pi-deprived (–Pi) seedlings of wild-type (WT), *pdr2*, and transgenic *p35S::LPR1* plants (OxL1). Seeds were germinated in +Pi liquid medium for 7 d prior to transfer to +Pi or –Pi media for 4 d. Whole root extracts were subjected to ammonium sulfate precipitation (40% saturation) and the supernatants (40 μL) were blotted and analyzed (anti-LPR1 antibodies). Membranes were stained with Coomassie-Blue.

**Extended Data Fig. 3.**
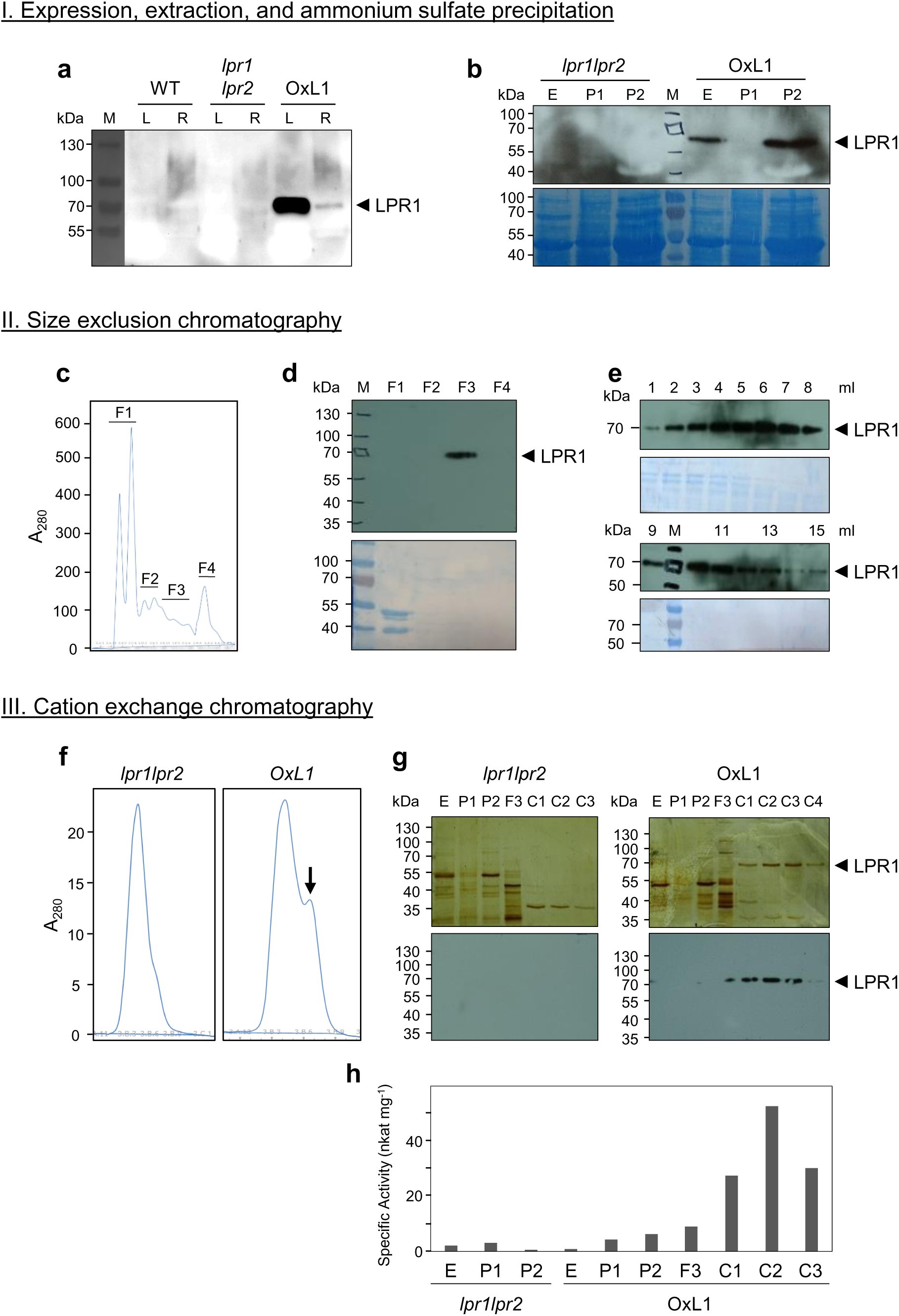
Three-step purification procedure of untagged native LPR1 protein to near homogeneity. **a.** Immunoblot analysis (n>10) of LPR1 protein expression in leaves (L) and roots (R) of wild-type (WT), *lpr1lpr2*, and transgenic *p35S::LPR1* (OxL1) plants (6-week-old, 50 μg total protein). **b.** Immunoblot analysis of protein extracts prepared from leaves of *p35S::LPR1* plants (E), and of protein pellets of fractions prepared by sequential ammonium sulfate precipitation: 40% saturation (P1) and 40-80% saturation (P2). Membrane stained with Coomassie-Blue. **c.** Elution profile after size-exclusion chromatography. Four major fractions (F1-F4) were pooled for further processing and analysis. **d.** Immunoblot analysis of pooled fractions (F1-F4). **e.** Immunoblot analysis of fraction F3 at 1-ml resolution. **f.** Elution profiles of protein preparations from *lpr1lpr2* and OxL1 plants after cation-exchange chromatography. Note the pronounced shoulder in the OxL1 profile (arrow). **g.** Separation of proteins by SDS-PAGE (upper panels: silver-stained gels,) and immunoblot analysis (lower panels) of all relevant fractions prepared from *lpr1lpr2* and OxL1 plants. **h.** Specific ferroxidase activities of the indicated fractions, (1µg total protein).

**Extended Data Fig. 4.**
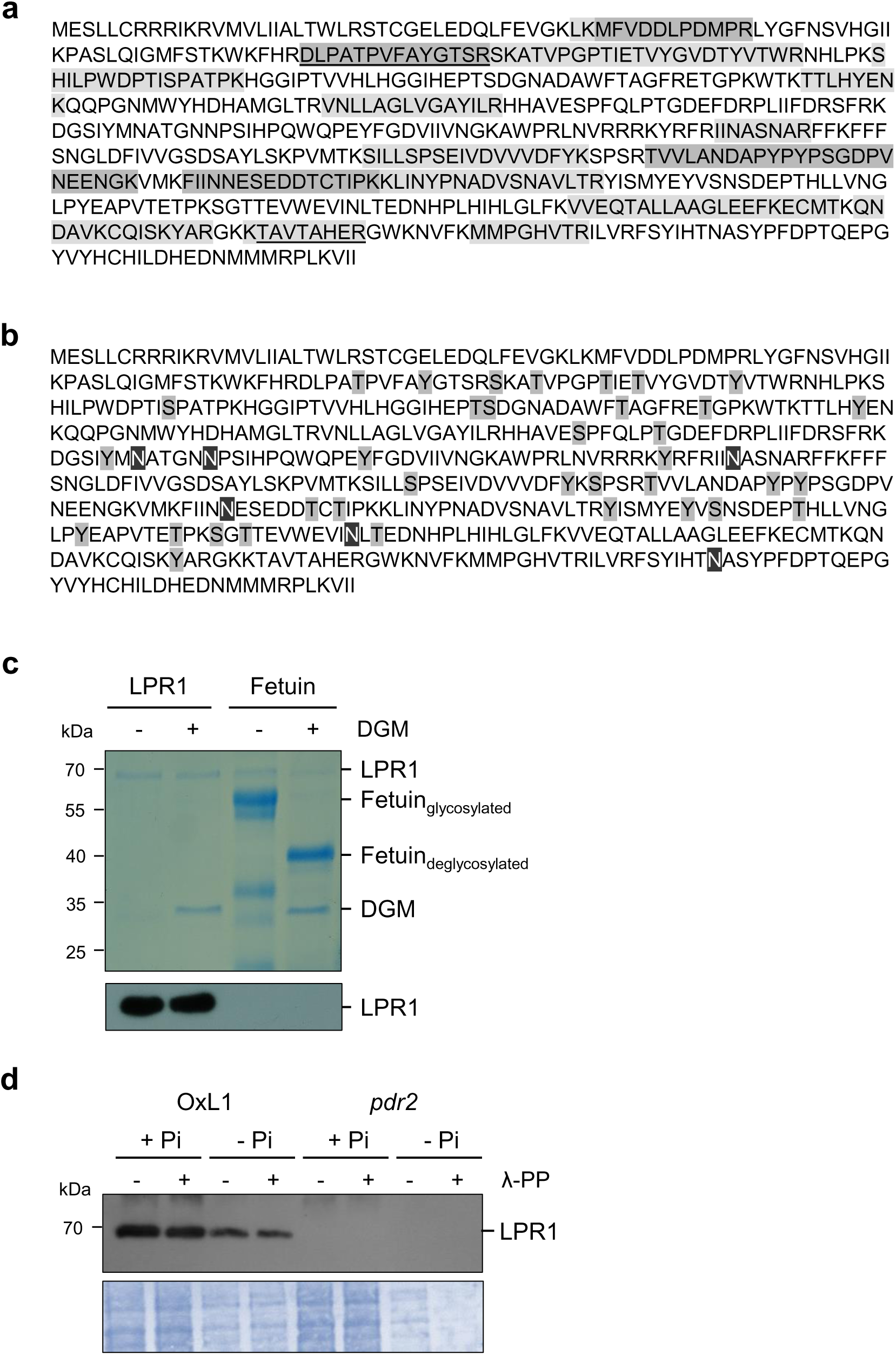
Verification and additional characterization of purified native LPR1. **a.** Native LPR1 protein purified from OxL1 leaf extracts was separated by SDS-PAGE. The identity of the eluted protein was determined by MS/MS peptide sequencing (n=5). Detected LPR1-derived peptides are highlighted (light grey) on the primary LPR1 structure; peptides detected in all five measurements are depicted in dark grey. Unique peptides identified in the TMT-dataset (quantitative proteomics) are underlined. **b.** Potential phosphorylation sites (PhosPhAt 4.0) are highlighted in grey and potential glycosylation sites are highlighted in black. **c.** Deglycosylation assay using purified LPR1 and fetuin protein as a positive control. Reactions with or without deglycosylating enzyme mix (DGM) were subjected to a 10% (w/v polyacrylamide) SDS-PAGE. Proteins were detected by Coomassie-Blue staining or immunoblot analysis with anti-LPR1 antibodies. **d.** Dephosphorylation assay using root extracts from OxL1 and *pdr2* plants germinated for 6 d on +Pi or –Pi media with and without Lambda protein phosphatase (n=2).

**Extended Data Fig. 5.**
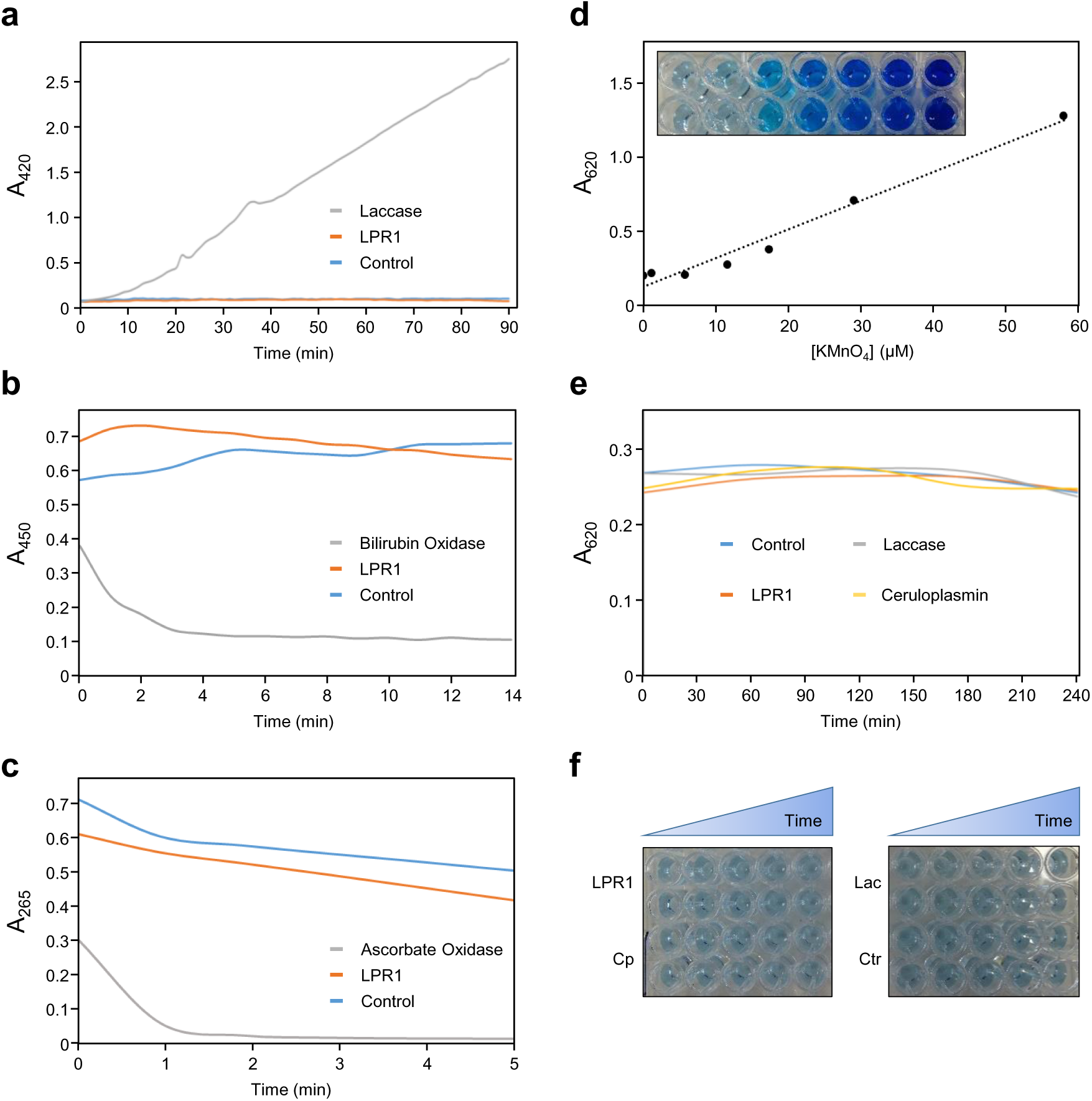
Substrate specificity of purified native LPR1. **a.** Laccase activity assay using 0.5 mM ABTS as the substrate and commercial laccase from *Trametes versicolor* as the control (n=4). **b.** Bilirubin oxidase assay using 35 μM bilirubin as the substrate and commercial bilirubin oxidase from *Myrothecium verrucaria* as the control (n=2). **c.** Ascorbate oxidase assay using 60 μM ascorbate as the substrate and commercial ascorbate oxidase from *Cucurbita* sp. as the control (n=4). **d-f.** Test for manganese oxidase activity using 1 mM MnSO_4_ as the substrate (n=3). Calibration curve of the KMnO_4_ (the oxidized product) in the presence of 0.005% (w/v) leucoberbelin blue (**d**). Assays with LPR1, laccase (*T. versicolor*) and human ceruloplasmin (ferroxidase), which did not display any detectable manganese oxidase activity (**e, f**).

**Extended Data Fig. 6.**
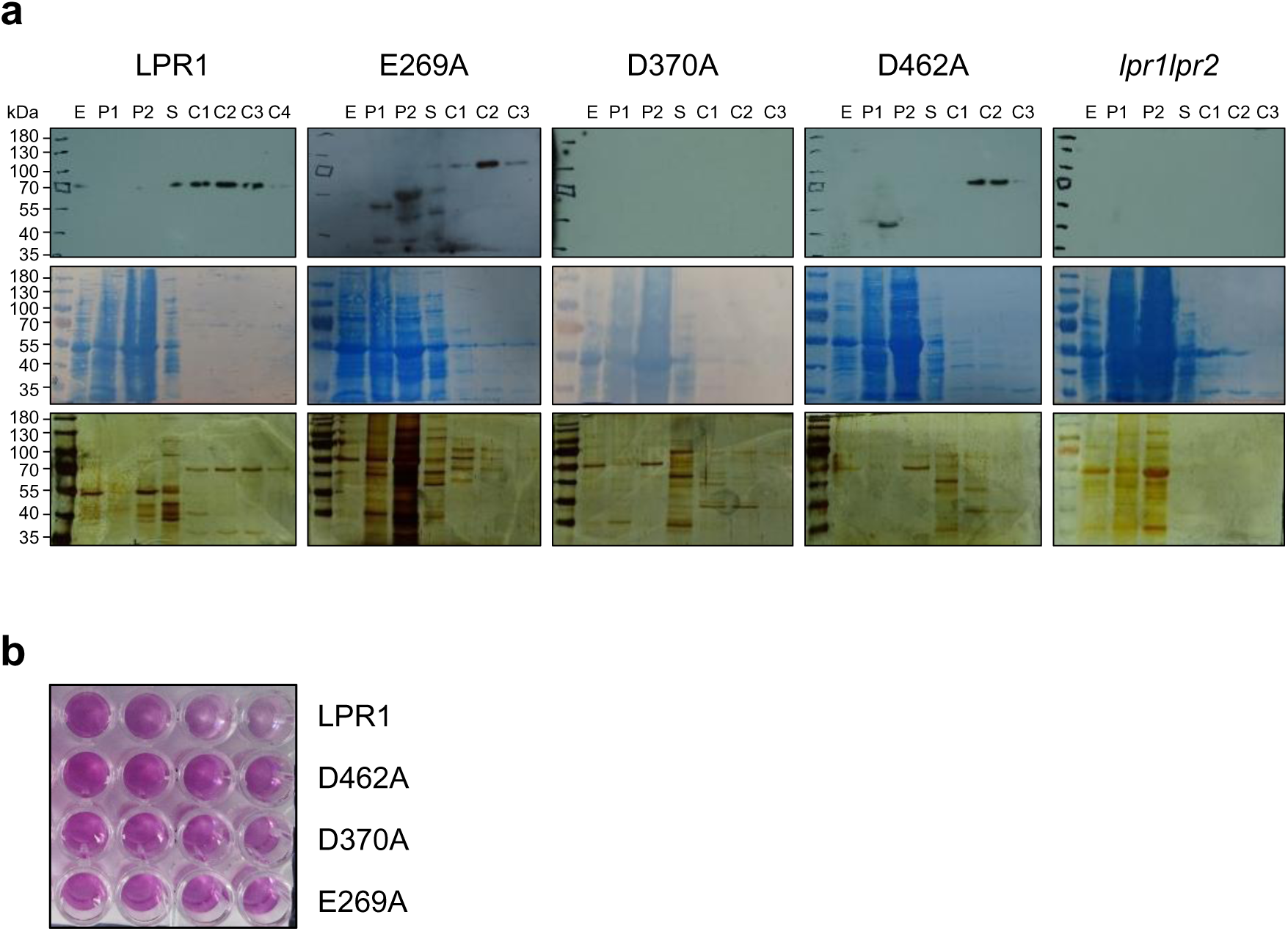
Purification of LPR1 protein variants. **a.** Using the three-step purification protocol, untagged LPR1 wild-type or mutant proteins were purified from extracts of transgenic *lpr1* plants expressing *p35S::LPR1* (LPR1), *p35S::LPR1^E269A^* (E269A), *p35S::LPR1^D370A^* (D370A) or *p35S::LPR1^D462A^* (D462A). Mutant line *lpr1lpr2* was used as negative control. Shown are immunoblots probed with anti-LPR1 antibodies (upper row), Coomassie Brilliant Blue-stained membranes of blots (center row), and silver-stained SDS gels loaded with 1 µg total protein per lane prior to separation (lower row). Protein fractions: (E) leaf extract; (P1) ammonium sulfate (40% saturation) precipitation; (P2) ammonium sulfate (40-80% saturation) precipitation; (S) size exclusion chromatography; (C1-C4) cation exchange chromatography. (n≥3) **b.** Discontinuous ferrozine assay using 1 µg of purified wild-type and mutant LPR proteins.

**Extended Data Fig. 7.**
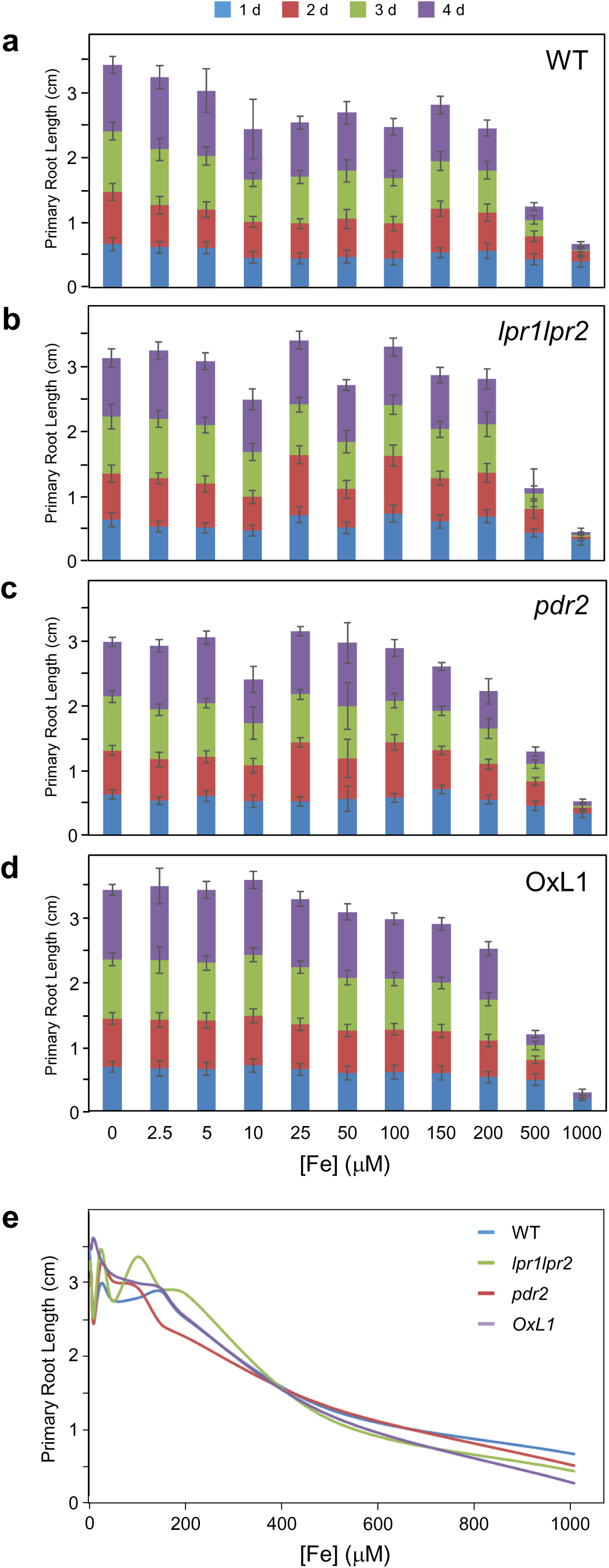
Fe-dependent primary root extension in Pi sufficiency. **a-d.** Seeds of wild-type (WT), l*pr1lpr2*, *pdr2*, and LPR1-overexpression (OxL1) plants were germinated for 5 d on +Pi agar prior to transfer to +Pi media with increasing iron (Fe^3+^-EDTA) supply. Gain of primary root extension was measured daily for up to 4 d after transfer and plotted for each genotype (±SD; n≥50). **e.** Trend lines of Fe-dependent primary root growth on +Pi media from 0-1000 μM Fe.

**Extended Data Fig. 8.**
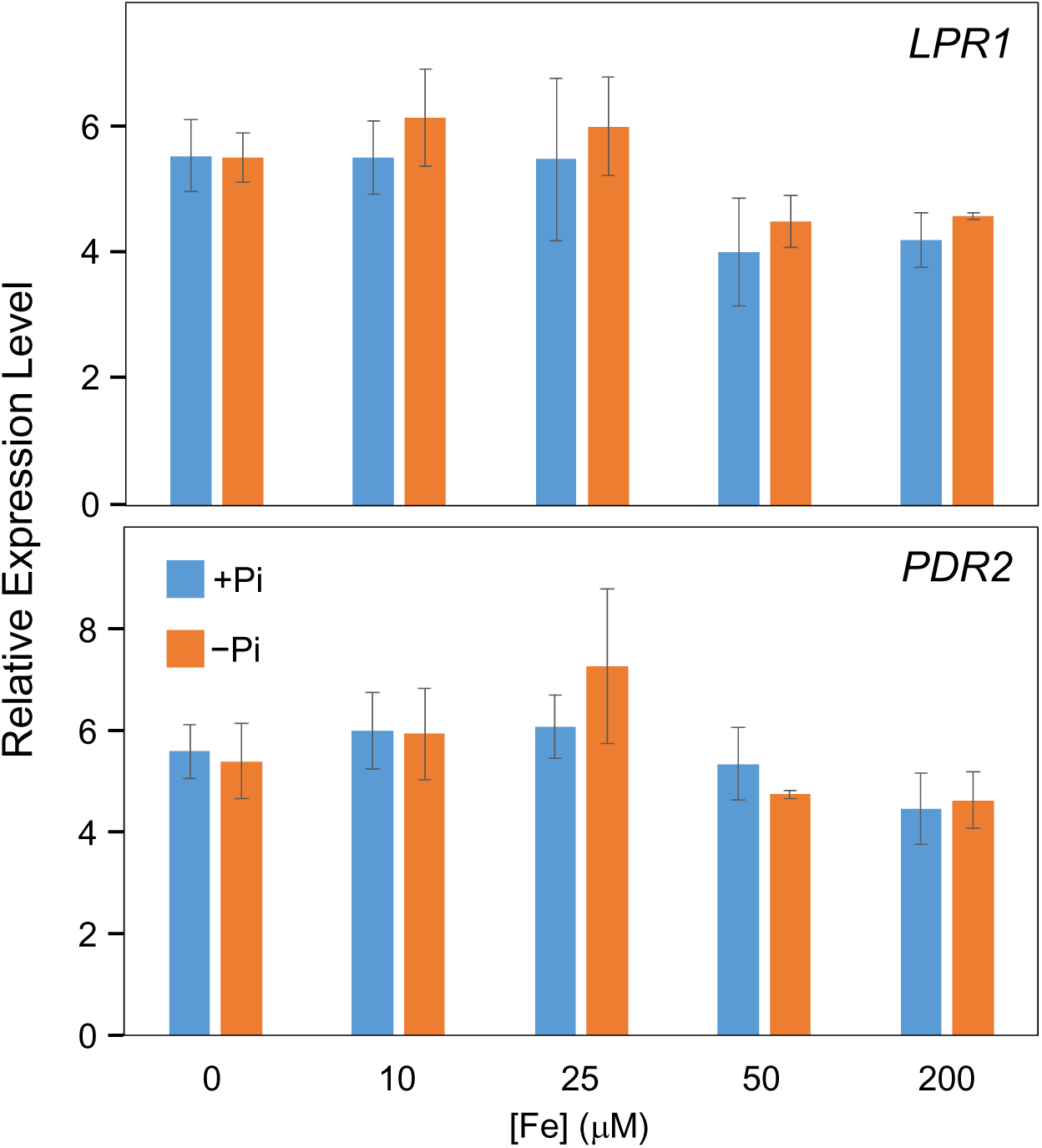
Fe-dependent regulation of *LPR1* and *PDR2* mRNA levels. Relative transcript levels (normalized to *UBC9* expression) of *LPR1* and *PDR2* in root tips. Wild-type (Col-0) seeds were germinated on +Pi medium (5 d) and transferred to +Pi or –Pi medium. After 24 h, the gain in root tip growth was harvested for RNA preparation and qRT-PCR analysis (± SD; n = 3).

**Extended Data Fig. 9.**
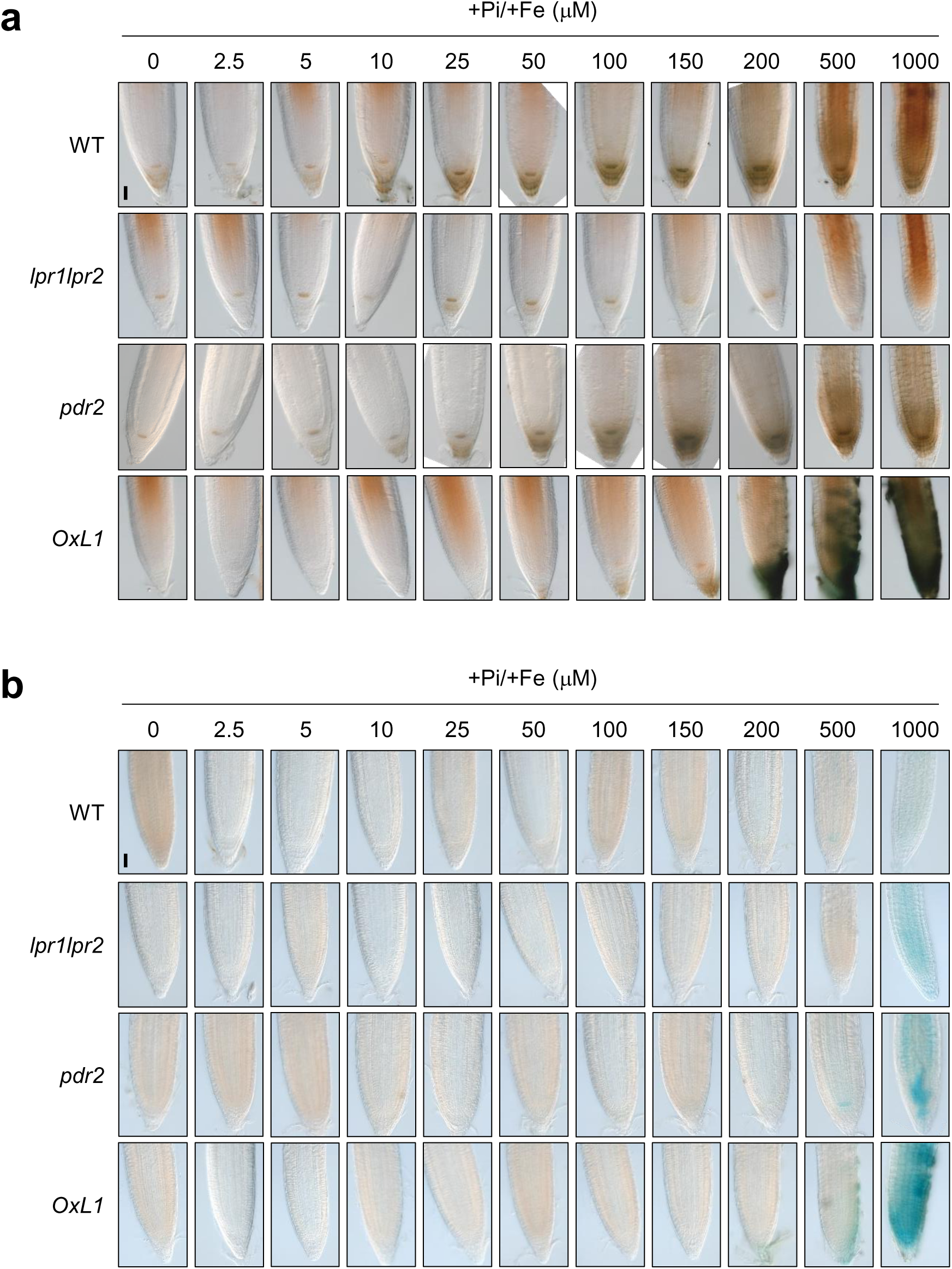
Fe-dependent Fe^3+^ accumulation in Pi-sufficient root tips. **a, b.** Seeds of wild-type (WT), *lpr1lpr2, pdr2* and LPR1-overexpression (OxL1) lines were germinated for 5 d on +Pi agar prior to transfer to +Pi media supplemented with increasing iron (Fe^3+^-EDTA) concentration. After 3 d of transfer, root tips were monitored for Fe^3+^ accumulation by Perls/DAB staining (**a**) and Perls staining only (**b**). Shown are representative images (n≥15). Scale bars 50µm.

**Extended Data Fig. 10.**
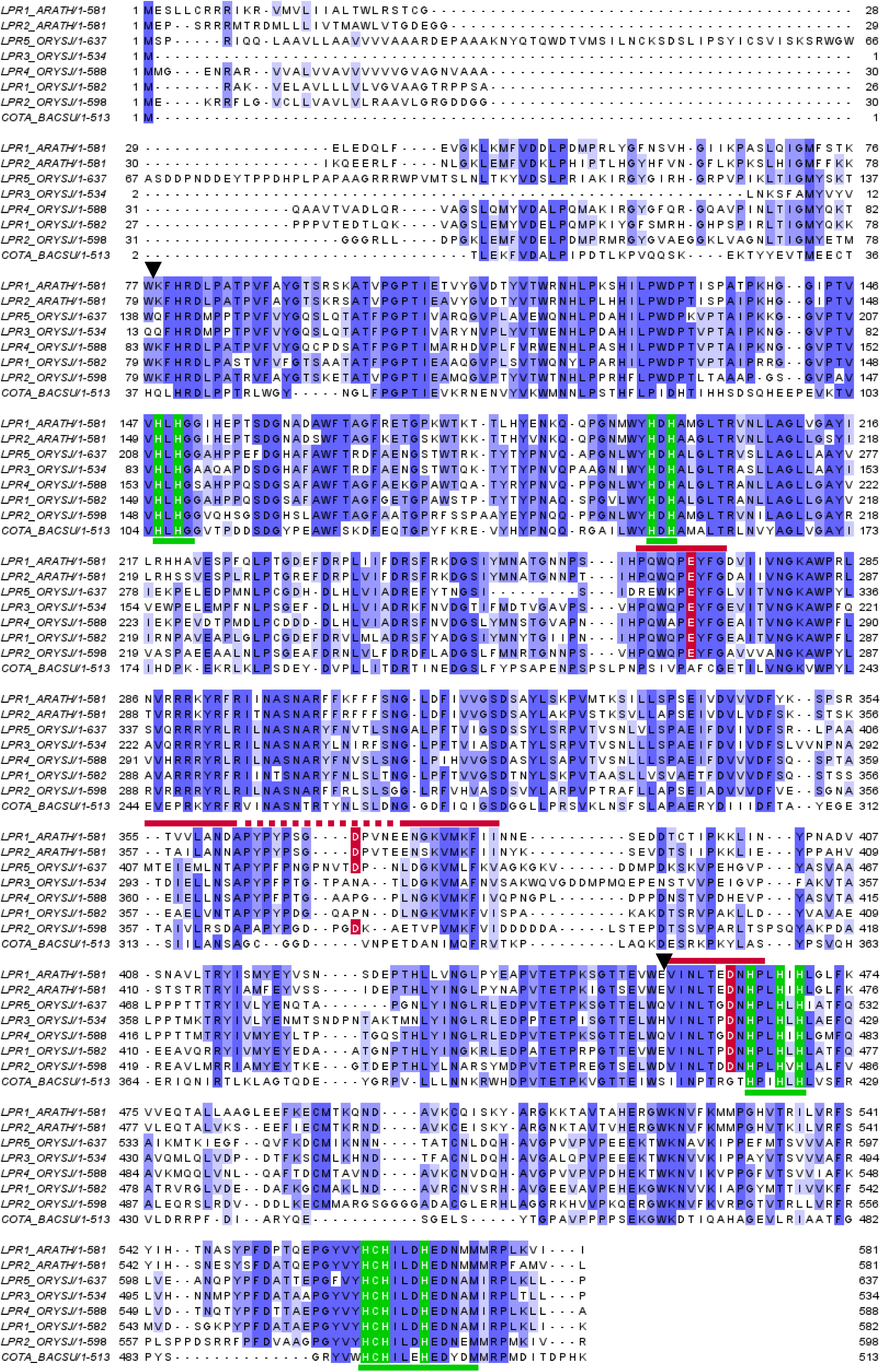
Primary sequence alignment of bacterial CotA and LPR1-related proteins from *Arabidopsis* and rice. Alignment of amino acid sequences of AtLPR1, AtLPR2, OsLPR1-5 and CotA (*Bacillus subtilis*). Shades of blue reflect degrees of positional sequence identity. Invariant histidine/cysteine residues of the T1 and T2/T3 copper cluster are shaded in green. The four conserved copper binding motifs typical for the MCO family (HXHG, HXH, HXXHXH, and HCHXXXHXXXXM/L/F) are delineated by green lines below the alignment. Residues of the acidic triad (presumed Fe^2+^ binding site) are depicted in red (E269, D370 and D462 on AtLPR1). Red lines above the alignment indicate conserved motifs flanking each residue of the acidic triad. Black triangles depict the position of conserved phase-0 introns.

**Extended Data Fig. 11.**
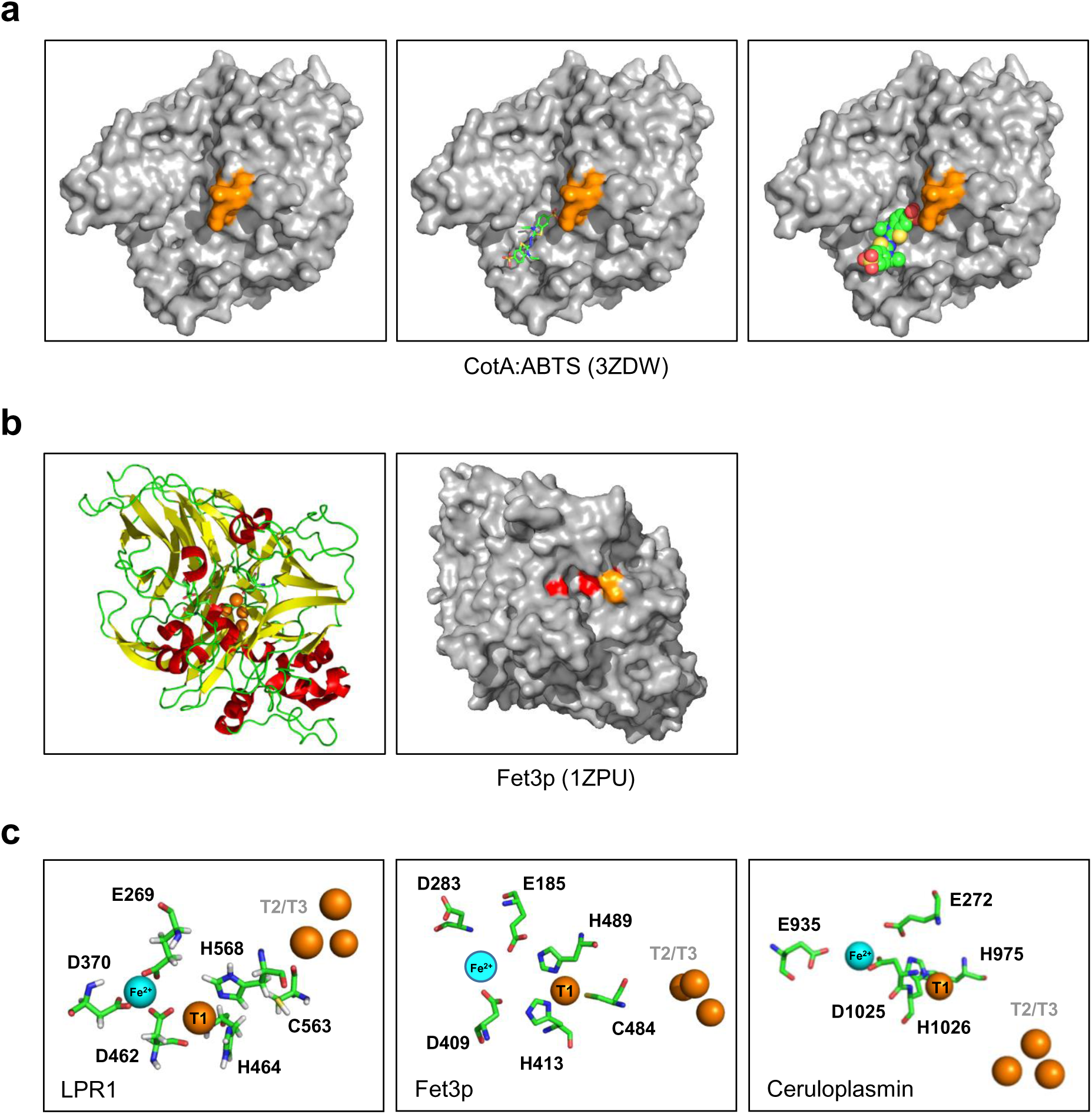
Substrate binding sites of CotA, LPR1, Fet3p, and ceruloplasmin. **a.** Surface representations of the CotA:ABTS complex (PDB ID: 3ZDW). The loop next to the substrate-binding site is highlighted (orange). Left panel: no ABTS. Center panel: ABTS (sticks). Right panel: ABTS (space filling). **b.** Experimental structure of Fet3p (PDB ID:1ZPU). Left panel: Ribbon presentation (Cu ions as orange spheres). Right panel: Surface representation (acidic triad in red, loop in orange). **c.** Structural models of the Fe^2+^ (blue sphere) binding site as well as T1 and T2/3 Cu (orange spheres) sites in LPR1, Fet3p, and ceruloplasmin (PDB ID: 1KCW).

**Extended Data Fig. 12.**
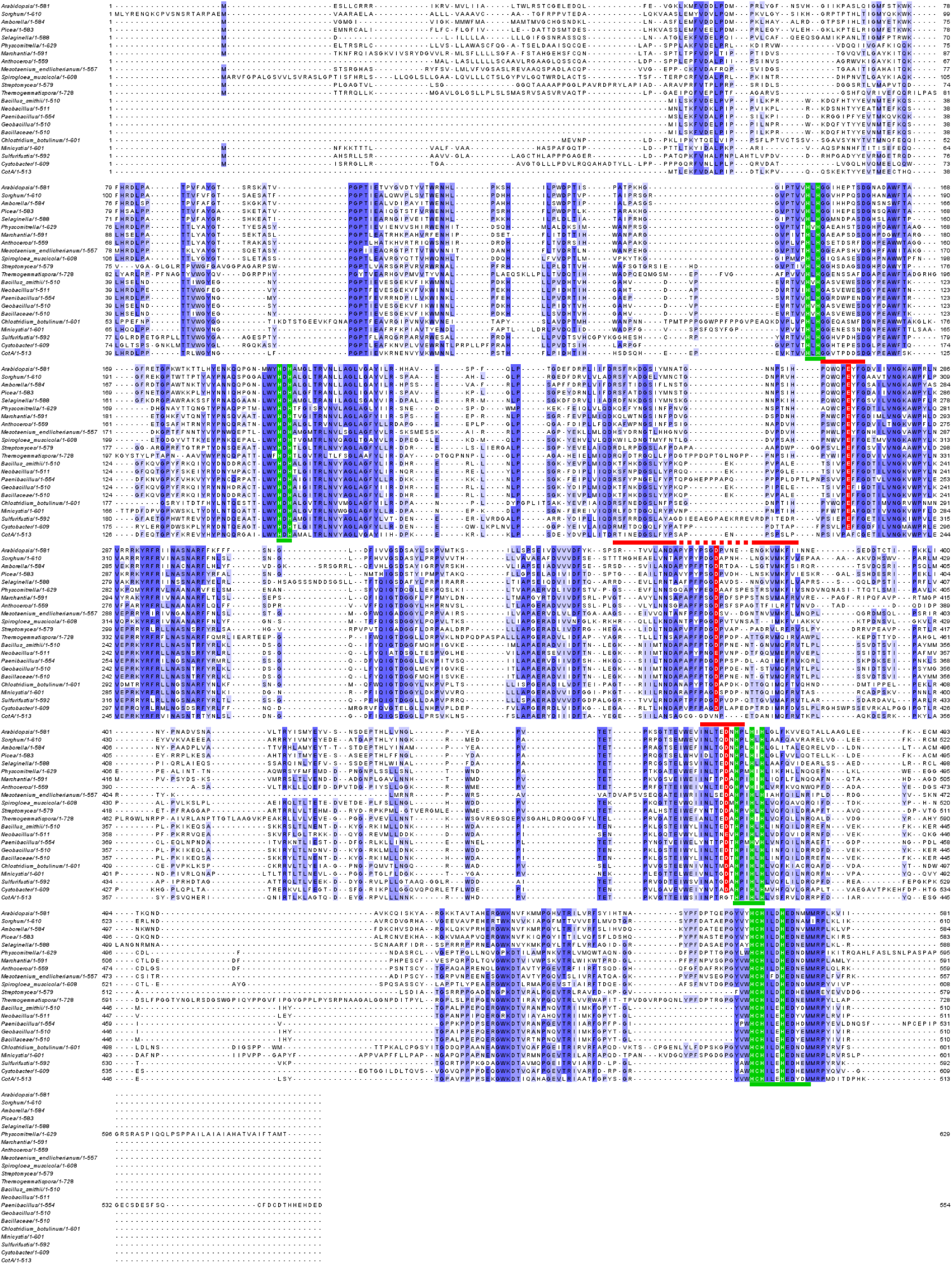
Primary sequence alignment of LPR1-like proteins from streptophytes and soil bacteria. Alignment of full-length amino acid sequences of *Arabidopsis* LPR1 and predicted LPR1-like proteins from select embryophytes, Zygnomatophyceae, and soil bacteria (see Fig. 3j). Shades of blue reflect degrees of positional sequence identity. Invariant histidine/cysteine residues of the T1 and T2/T3 copper cluster are shaded in green. The four conserved copper-binding motifs typical for the MCO family (HXHG, HXH, HXXHXH, and HCHXXXHXXXXM/L/F) are delineated by green lines below the alignment. Residues of the acidic triad (presumed Fe^2+^ binding site) are depicted in red (E269, D370 and E462 on AtLPR1). Red lines above the alignment indicate conserved motifs flanking each residue of the acidic triad.

**Extended Data Fig. 13.**
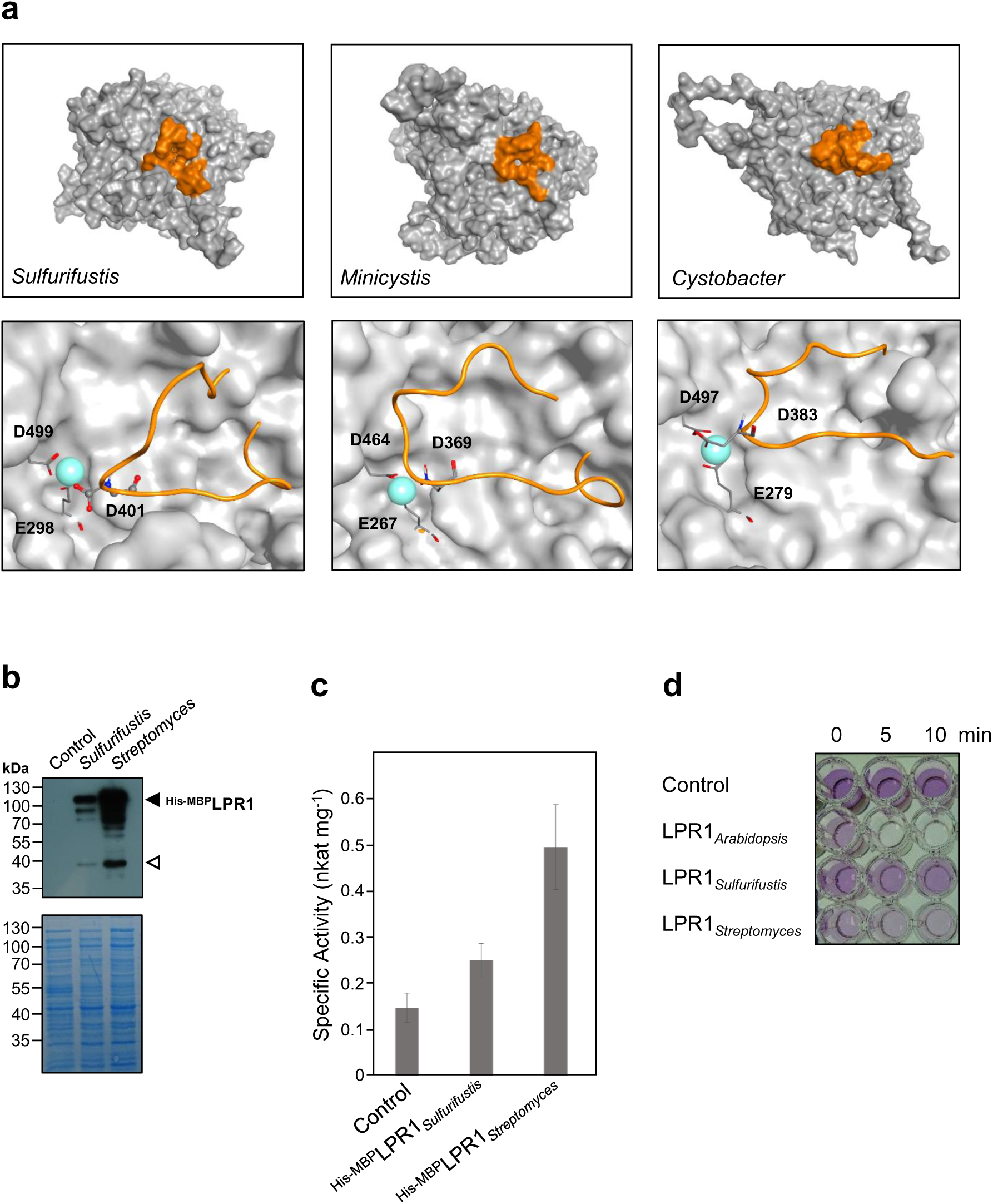
Bacterial LPR1-like proteins reveal ferroxidase activity. **a.** Homology modeling of LPR1-like proteins from *Sulfurifustis variabilis*, *Minicystis rosea* and *Cystobacter fuscus* using CotA as the template. The top panels show surface representations with the linker sequence harboring the central residue of the acidic triad highlighted in orange. The enlargements (lower panels) show the loop in ribbon (orange), the residues of the acidic triad as sticks, and the Fe^2+^ substrate cation as blue sphere. **b.** Expression of affinity-tagged LPR1-like proteins from *Sulfurifustis* and *Streptomyces* in *E. coli* (n=4). **c.** Specific ferroxidase activity of *E. coli* extracts after induction of recombinant protein expression (n=3). **d.** Ferrozine microtiter plate assay for ferroxidase activity with *E. coli* extracts.

**Extended Data Fig. 14.**
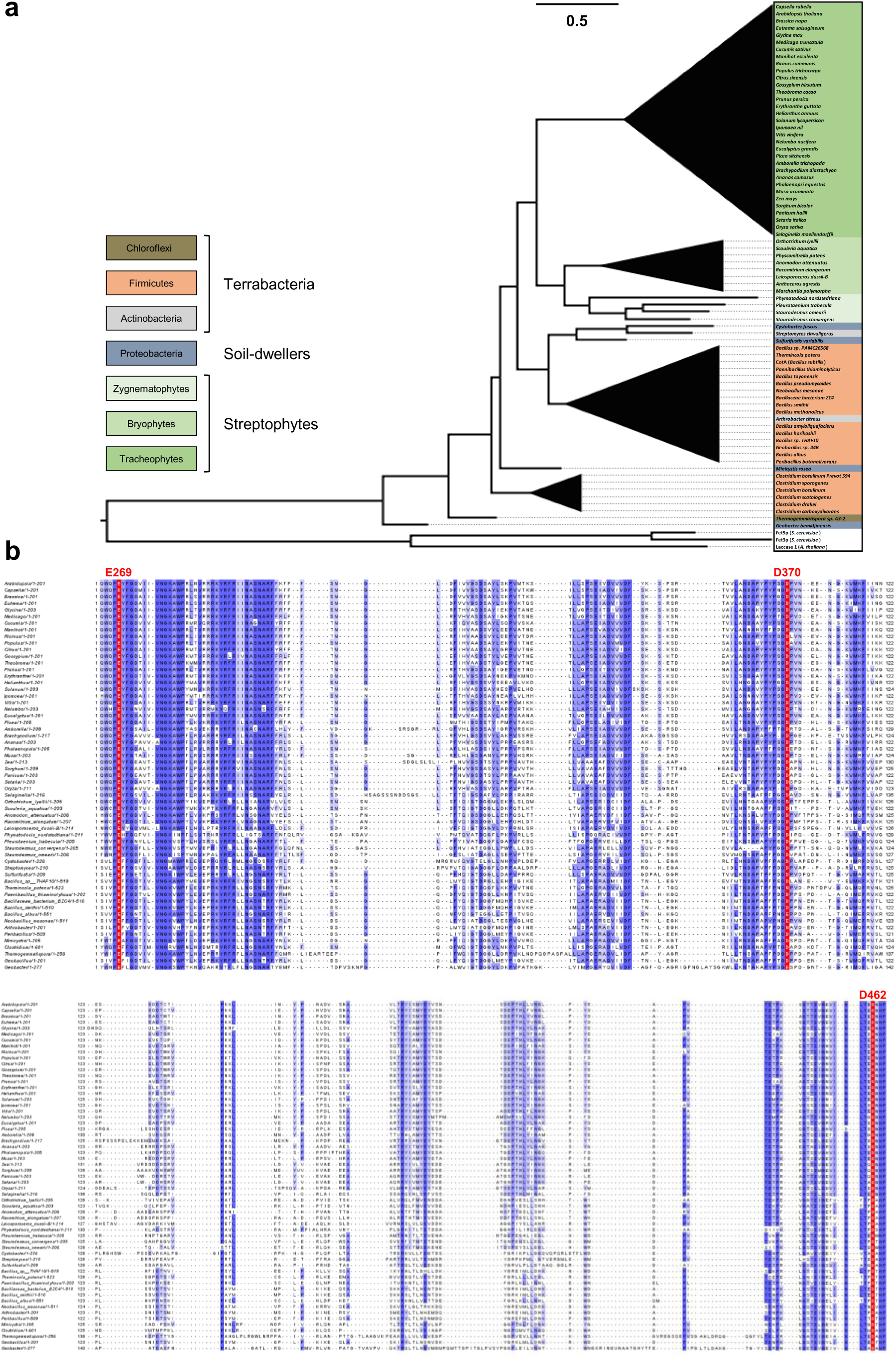
Phylogenetic relationship of LPR1-like MCO proteins. **a.** Phylogenetic tree of predicted LPR1-like MCO amino acid sequences from bacteria, Zygnematophyceae and embryophytes (see Extended Data Table 2). Full-length bacterial coding DNA sequences (CDS) and streptophyte CDS starting from the predicted second intron (see Fig 8b) were conceptually translated and used to generate the maximum-likelihood midpoint-rooted tree (450 bootstrap replicates). Major groups are collapsed (triangles). **b.** Alignment of polypeptide sequences that contiguously cover the three motifs of the acidic triad (predicted Fe-binding site) of LPR1-like proteins relative to *Arabidopsis* LPR1 (E269, D370, and D462, red shade). Shades of blue reflect degrees of positional sequence identity.

**Extended Data Fig. 15.**
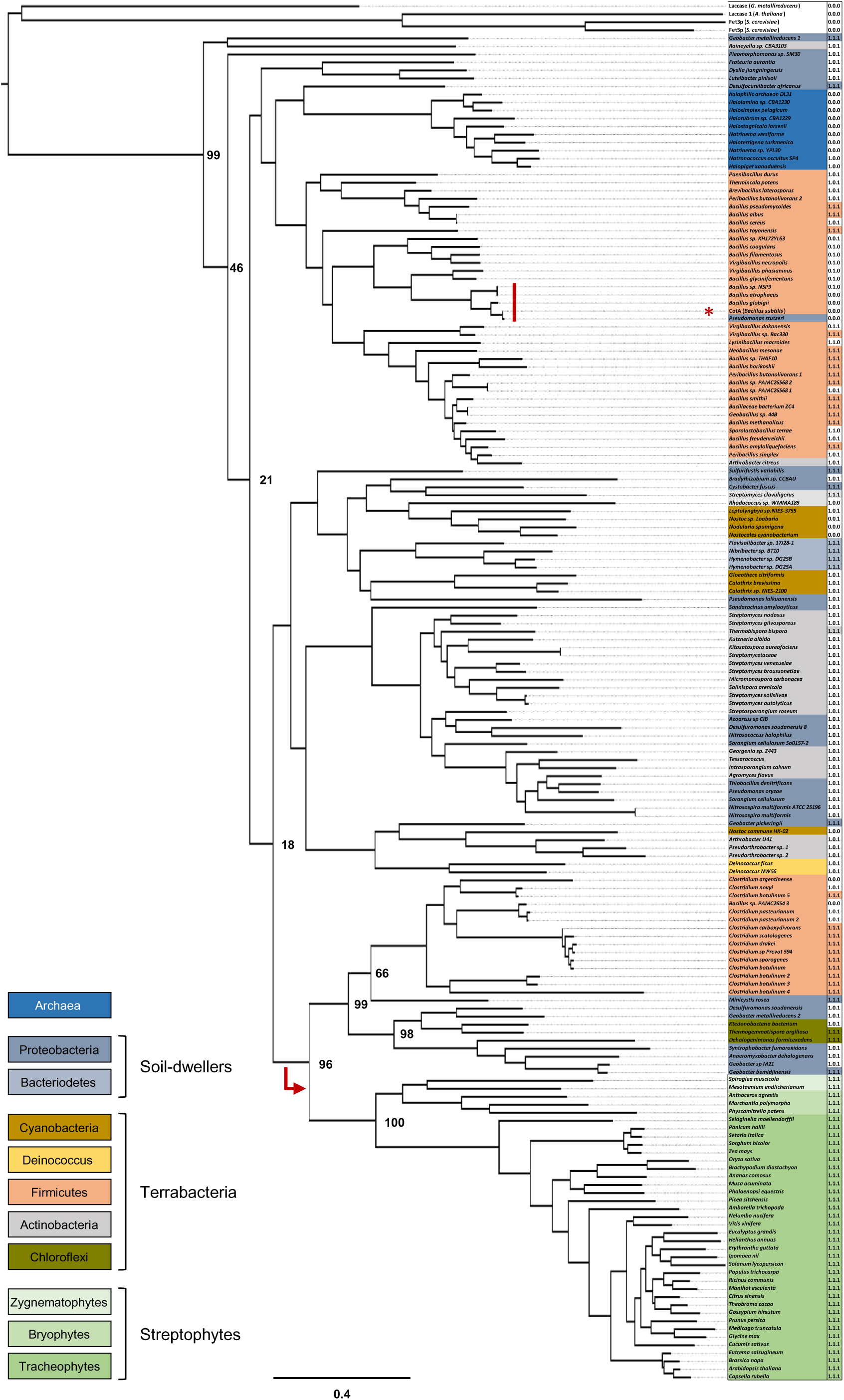
Evolution of LPR1-type MCO proteins. Phylogenetic tree of predicted LPR1-like MCO amino acid sequences from bacteria, Zygnematophyceae and embryophytes (see Extended Data Table 3). Bacterial and streptophyte coding DNA sequences were conceptually translated and used to generate the maximum-likelihood midpoint-rooted tree (350 bootstrap replicates). The presence of an acidic triad motif is indicated by numerical code on the right of the scientific name: complete (1.1.1; colored), partial (1.0.1 and combinations) or absent (0.0.0). CotA (*B. subtilis*) and highly similar proteins are indicated by a red asterisk and vertical bar, respectively. The streptophyte sequences occupy a monophyletic clade nested within the paraphyletic bacterial radiation, which suggests a single horizontal gene transfer event from a bacterial donor (red arrow).

**Extended Data Table 1.**
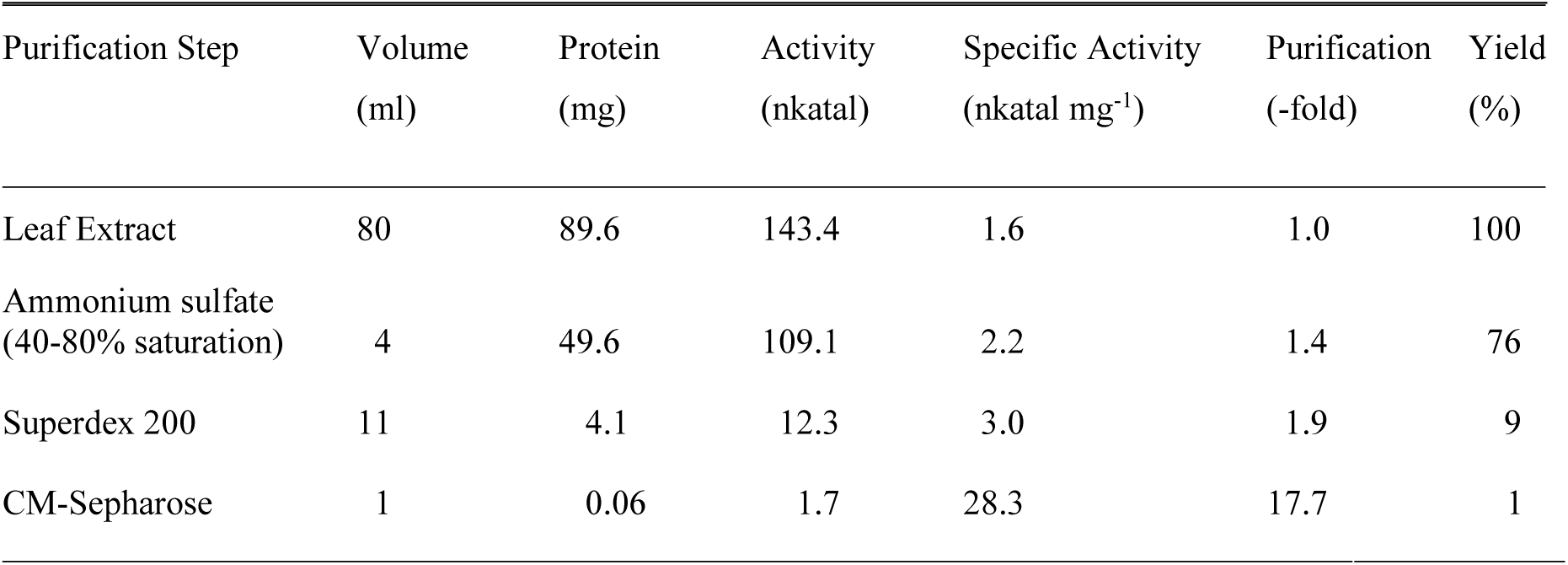
Purification of native *Arabidopsis* LPR1 protein. Native, untagged LPR1 protein was purified from leaves of transgenic *p35S::LPR1* plants (*A. thaliana*). All fractions were assayed for ferroxidase activity at pH 5.8 as described.

**Extended Data Table 2.**
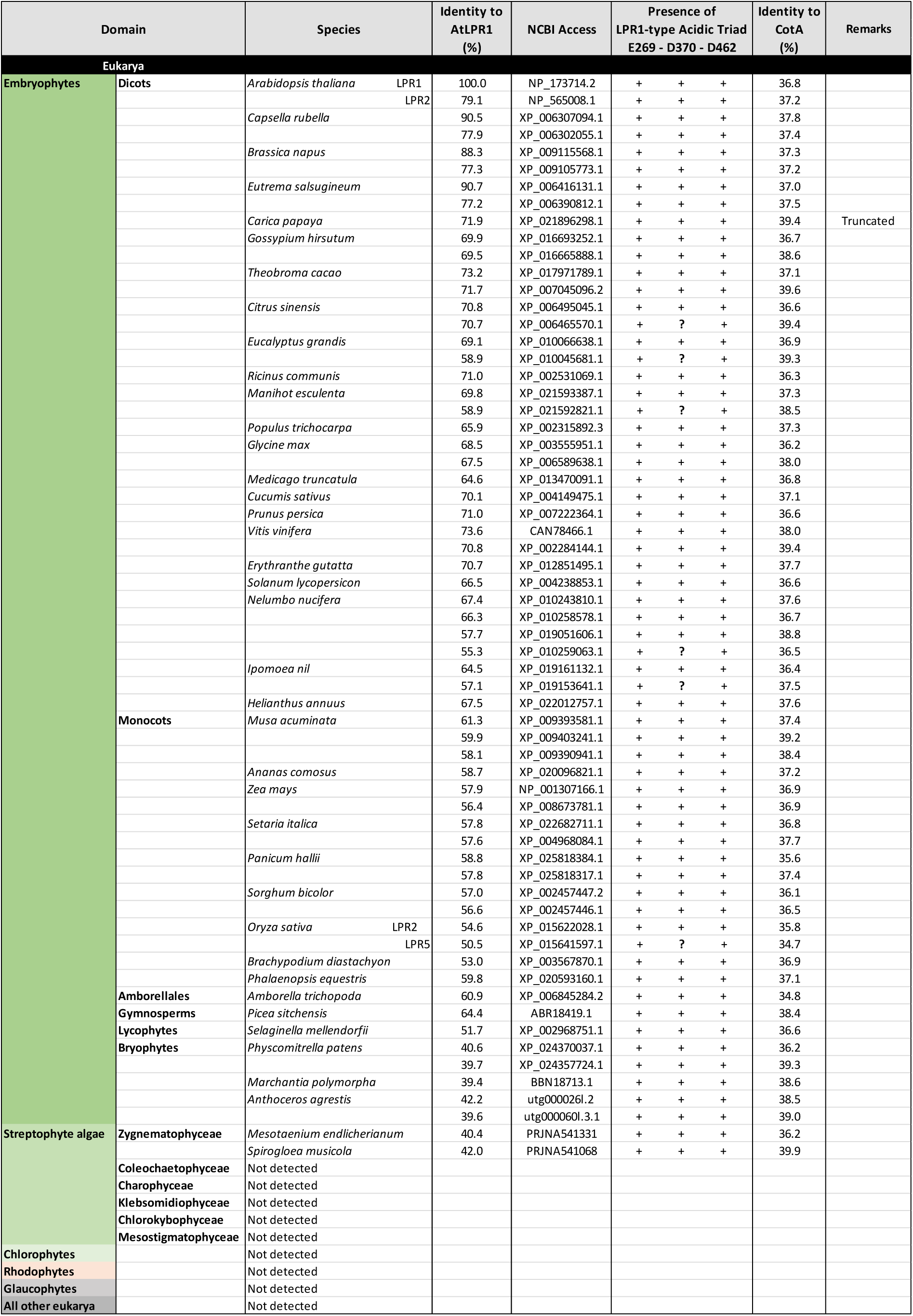
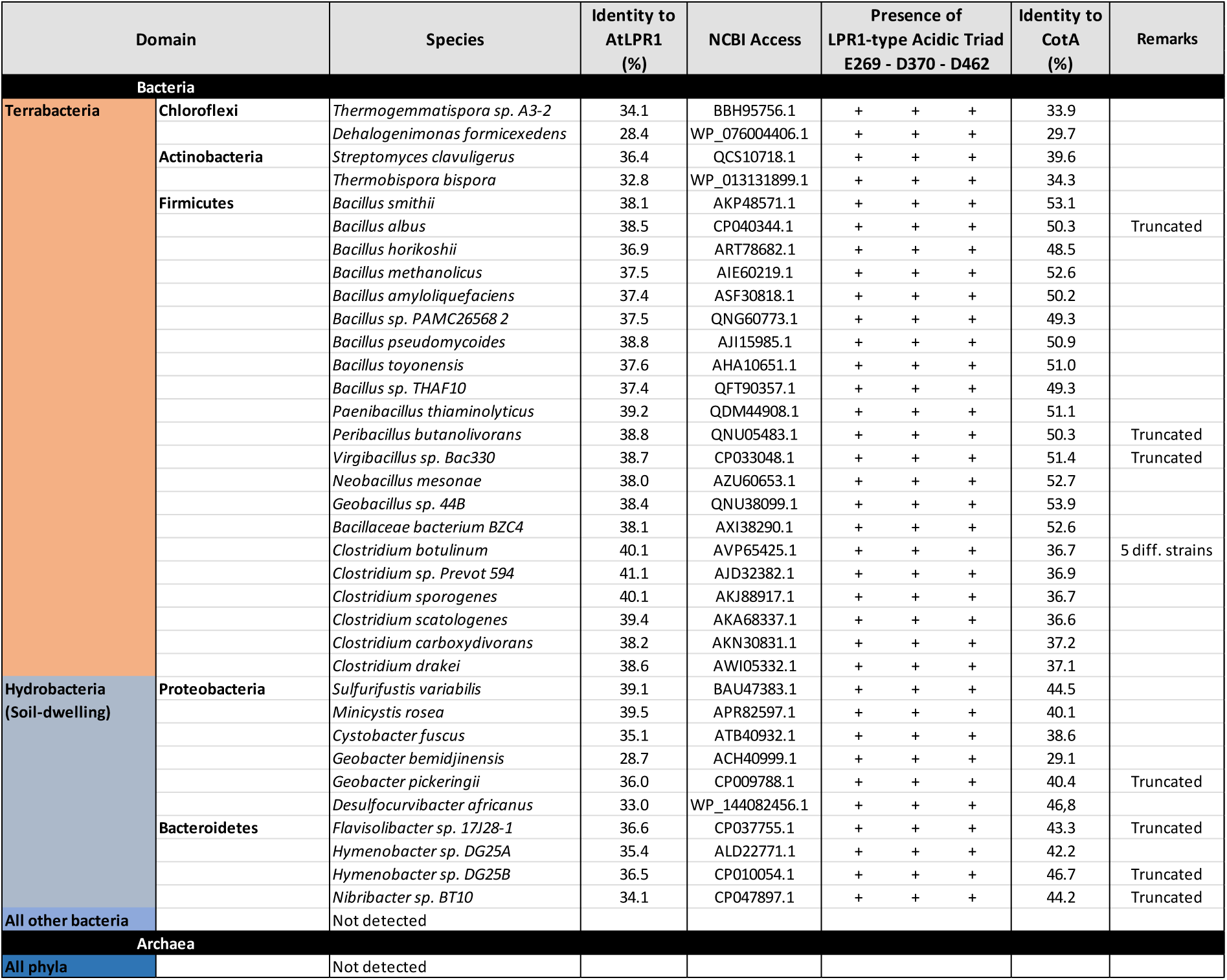
Distribution of LPR1-type MCO ferroxidases across the domains of life. Embryophyte LPR1-like sequences were identified by BLASTP searches at NCBI using *Arabidopsis thaliana* LPR1 (At1g23010) as the query. Hits for select land plant species are listed. A LPR1 profile HMM (14 species) was used to interrogate at NCBI all other domains and phyla. Hits were filtered and visually inspected to validate the presence of conserved acidic triad motifs necessary for Fe-binding by LPR1 (see Extended Data Table 3). The same HHM approach was used to search transcript data of the 1KP Project (One Thousand Plant Transcriptomes Initiative 2019) and the recently published genomes of Anthoceros angustus (Li et al. 2020), *Mesotaenium endlicherianum* and *Spirogloea musicola* (Cheng et al. 2019), and *Mesostigma viride* and *Chlorokybus atmophyticus* (Wang et al. 2020). One Thousand Plant Transcriptomes Initiative (2019) Nature 574:679-695 Li et al. (2020) Nat Plants 6:259-272 Cheng et al. (2019) Cell 179:1057-1067 Wang et al. (2020) Nat Plants 6:95-106

**Extended Data Table 3.**
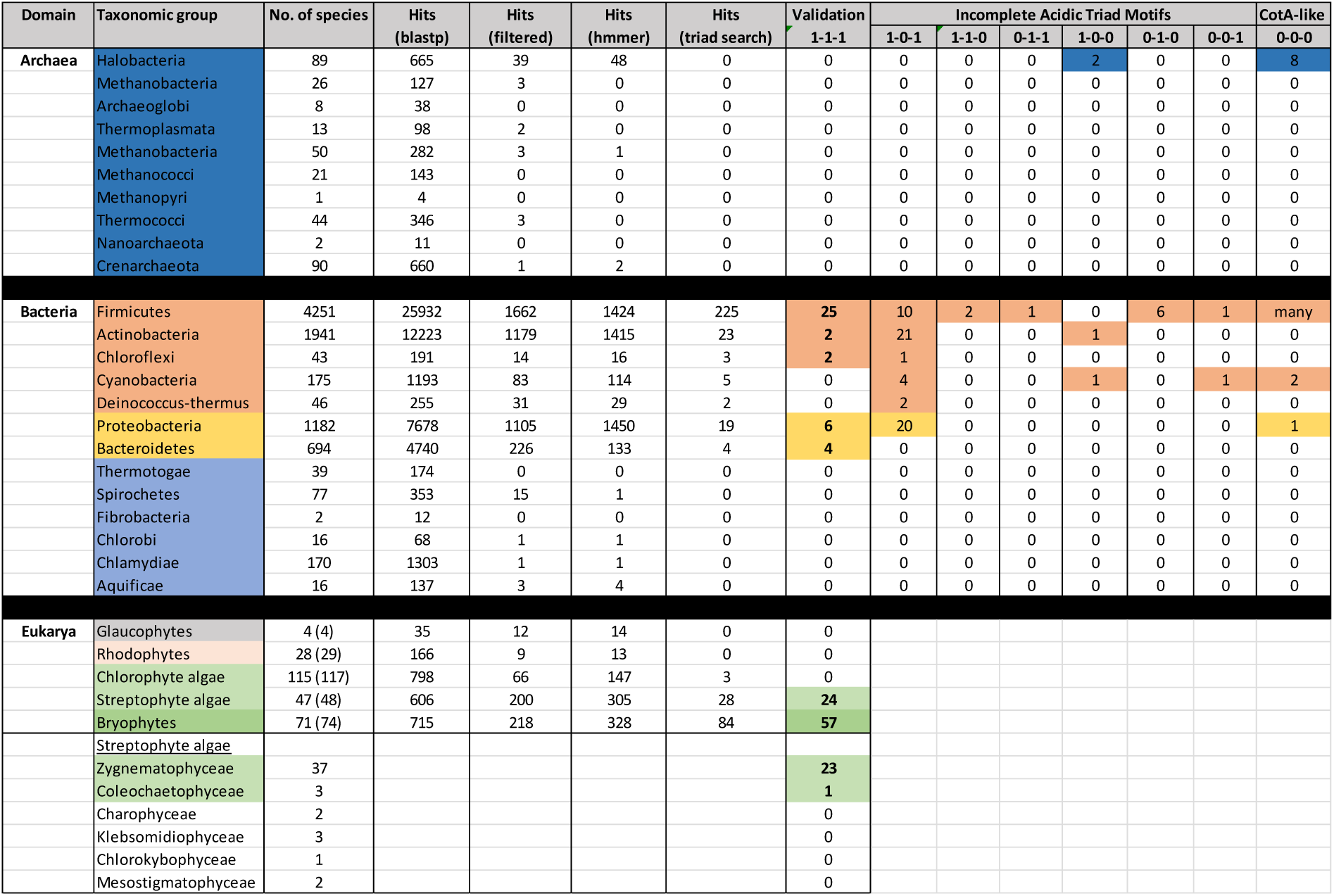
HMM profile search for sequences related to LPR1-type MCO ferroxidases. A LPR1 profile HMM (14 species) approach was used to search the domains Bacteria and Archaea as well as the transcript data of the 1KP Project (One Thousand Plant Transcriptome Initiative 2019). Hits were filtered and visually inspected to validate the presence of conserved acidic triad motifs around E269, D370 and D462 (abbreviated as 1-1-1) necessary for Fe-binding by LPR1 (see Extended Data Table 2). Additional searches at NCBI were conducted, using the internal polypeptide of LPR1 covering the entire acidic triad segment (aa 264-465) as well as of CotA (aa 222-420) as the queries, to identify additional LPR1-related MCO sequences with incomplete acid triad motifs (abbreviated as 1-0-1 to 0-0-0). One Thousand Plant Transcriptomes Initiative (2019) Nature 574:679-695

**Extended Data Table 4.**
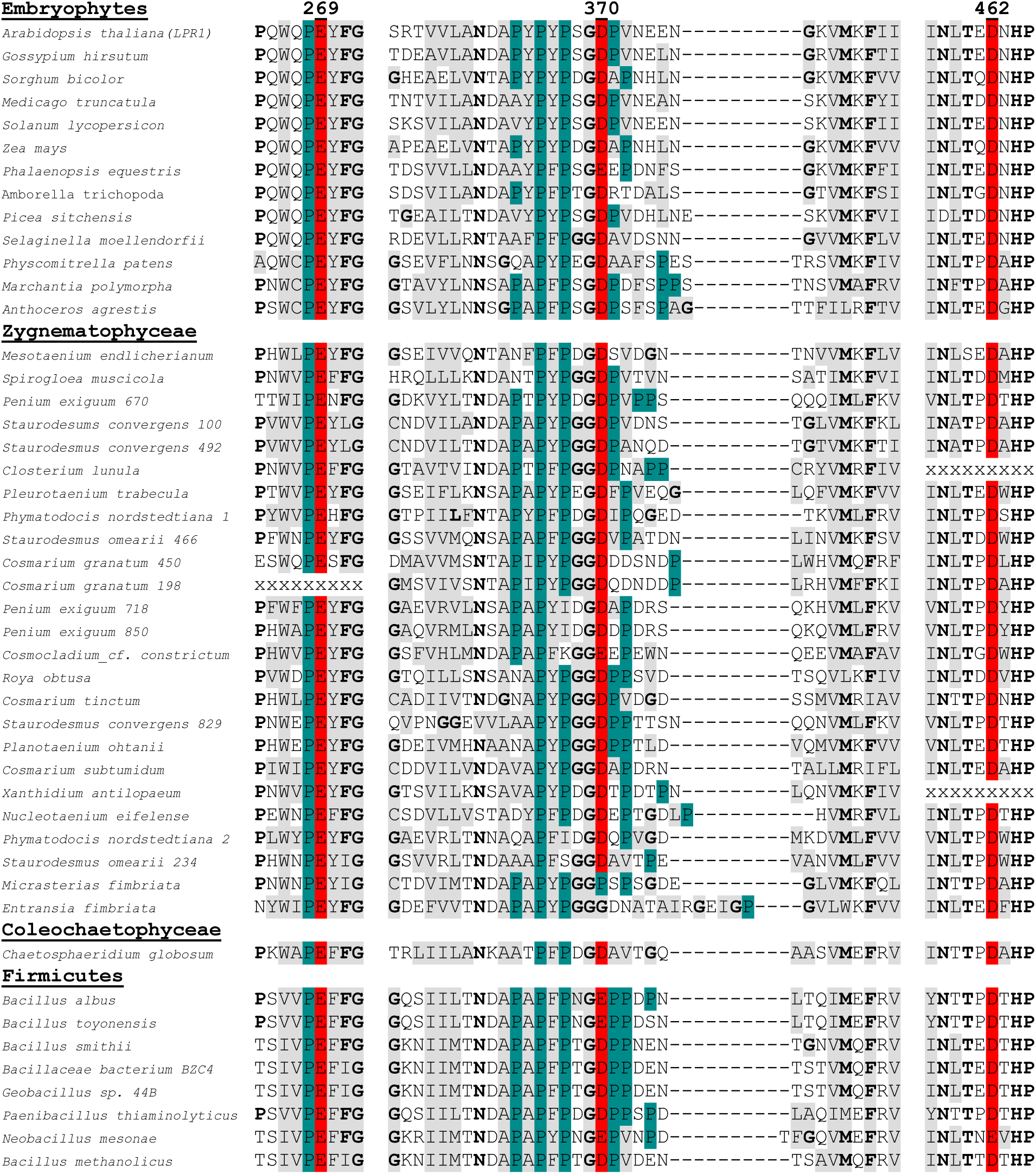
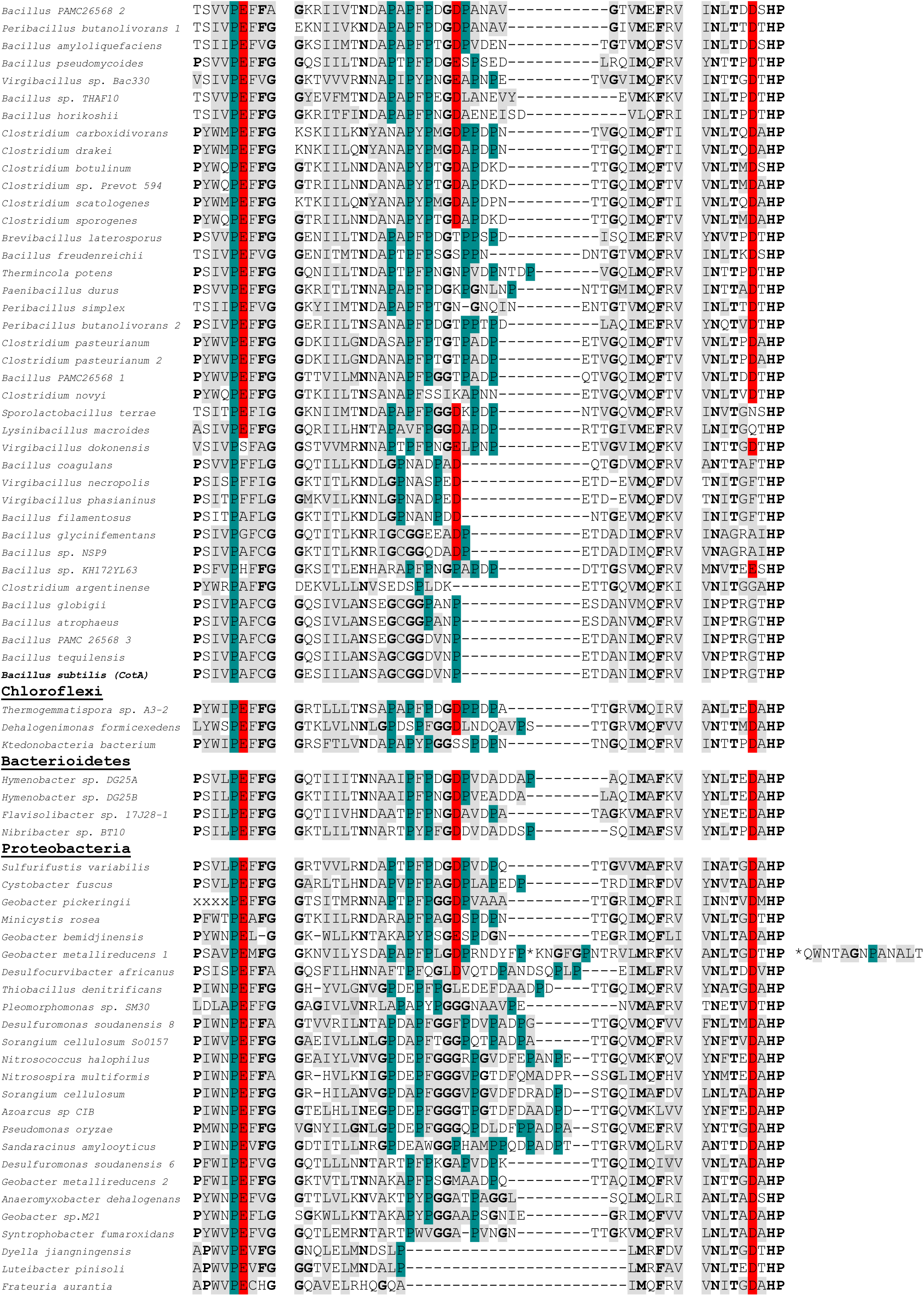
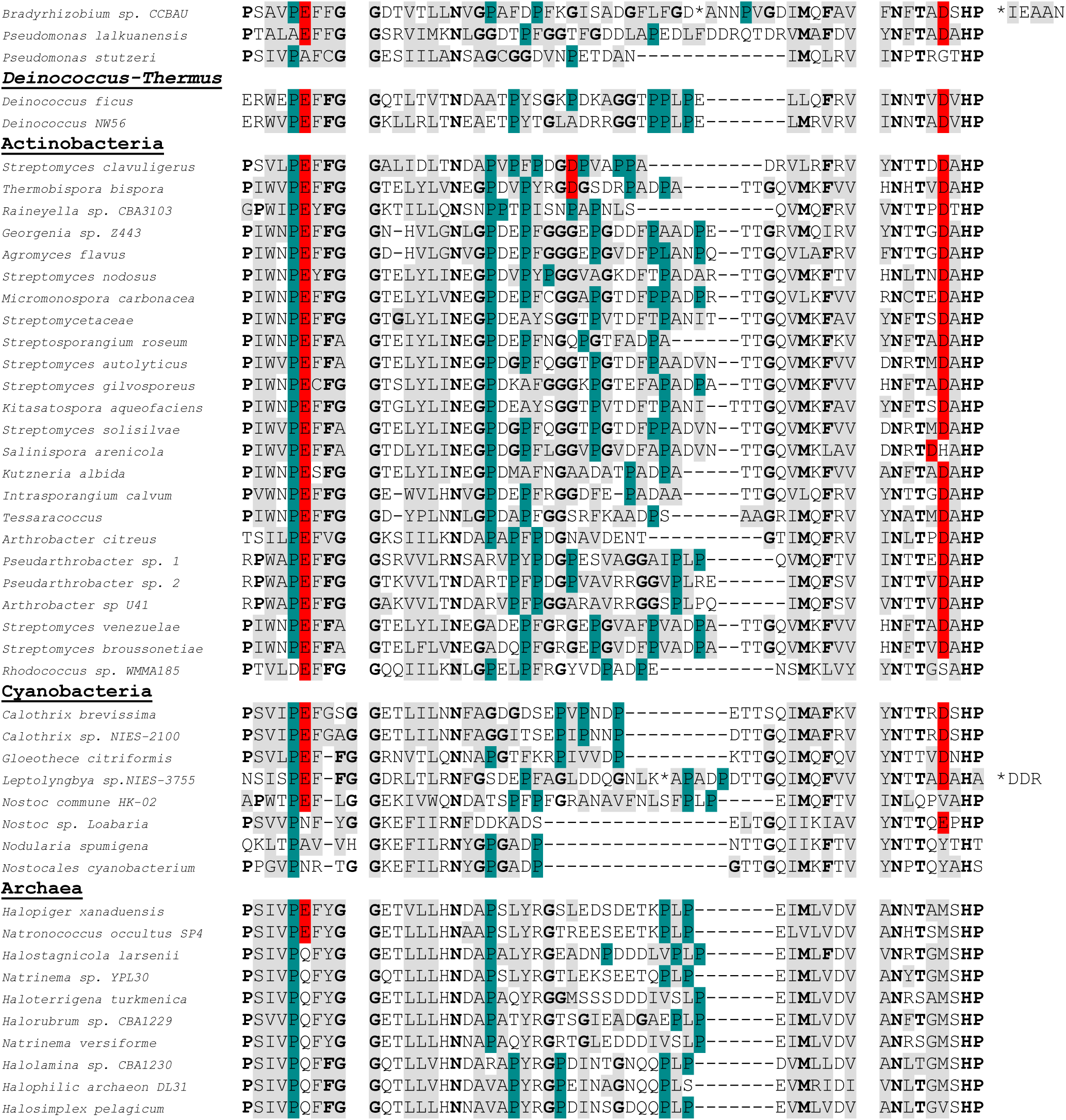
Alignment of LPR1-related acidic triad motif polypeptide sequences. Exhaustive tblastn searches were conducted (NCBI nucleotide collection; One Thousand Plant Transcriptomes Initiative 2019, known as 1KP Project) to identify polypeptide sequences related to the conserved motifs flanking the acidic triad residues of Arabidopsis LPR1 ferroxidase. All sequences of are aligned relative to E269, D370, and D462 of Arabidopsis LPR1 (highlighted in red). Hydrophobic amino acid residues are highlighted in grey. Proline residues of the first motif (E269), except at the start, and in the center of the variable linker (flanking D370) are highlighted in petrol blue. Glycine residues in the variable linker segment, and highly frequent or invariant residues in all three motifs are typed in bold face. Aligned are the sequences of select embryophytes (see Extended Data Table 2), identified in streptophyte algae by the 1KP Project, and retrieved from NCBI for bacterial phyla. Note, sequences for some streptophyte algae are truncated (xxx). Three sequences are too long for reasonable alignment (*, extra peptide sequences on the right).

